# Designing antimicrobials with programmable mechanism and safety

**DOI:** 10.64898/2026.09.01.747572

**Authors:** Paulina Szymczak, Marcelo Der Torossian Torres, Diogo Soares, Leon Hetzel, Bruno Puczko-Szymański, Stefanie Jegelka, Stephan Günnemann, Fabian J. Theis, Cesar de la Fuente-Nunez, Ewa Szczurek

## Abstract

Antimicrobial peptides (AMPs) are a promising solution to antimicrobial resistance, yet generative models for their design cannot control the physicochemical properties and motifs that shape activity and selectivity. Here, we present OmegAMP, a conditional diffusion framework controlling net charge, mean hydrophobicity, and sequence length, supporting *de novo*, analog, and motif-guided design. Across 204 wet-lab characterized peptides, *de novo* generation yielded antimicrobials with broad activity against multidrug-resistant Gram-negative isolates. Analog generation converted six inactive prototypes into antimicrobials, with the prototype determining each analog’s membrane-disruption mode and mammalian-cell safety. Motif-guided analog generation preserved lipopolysaccharide engagement of active prototypes, and a redesigned non-antimicrobial leucine zipper acquired antimicrobial activity while retaining DNA-perturbing character *in vitro*. In murine skin and thigh infection models, leads reduced bacterial burden, with a motif-guided DNA-perturbing lead matching the fluoroquinolone control systemically. OmegAMP opens a programmable route to new peptide antibiotics whose mechanism and safety follow from the chosen prototype.

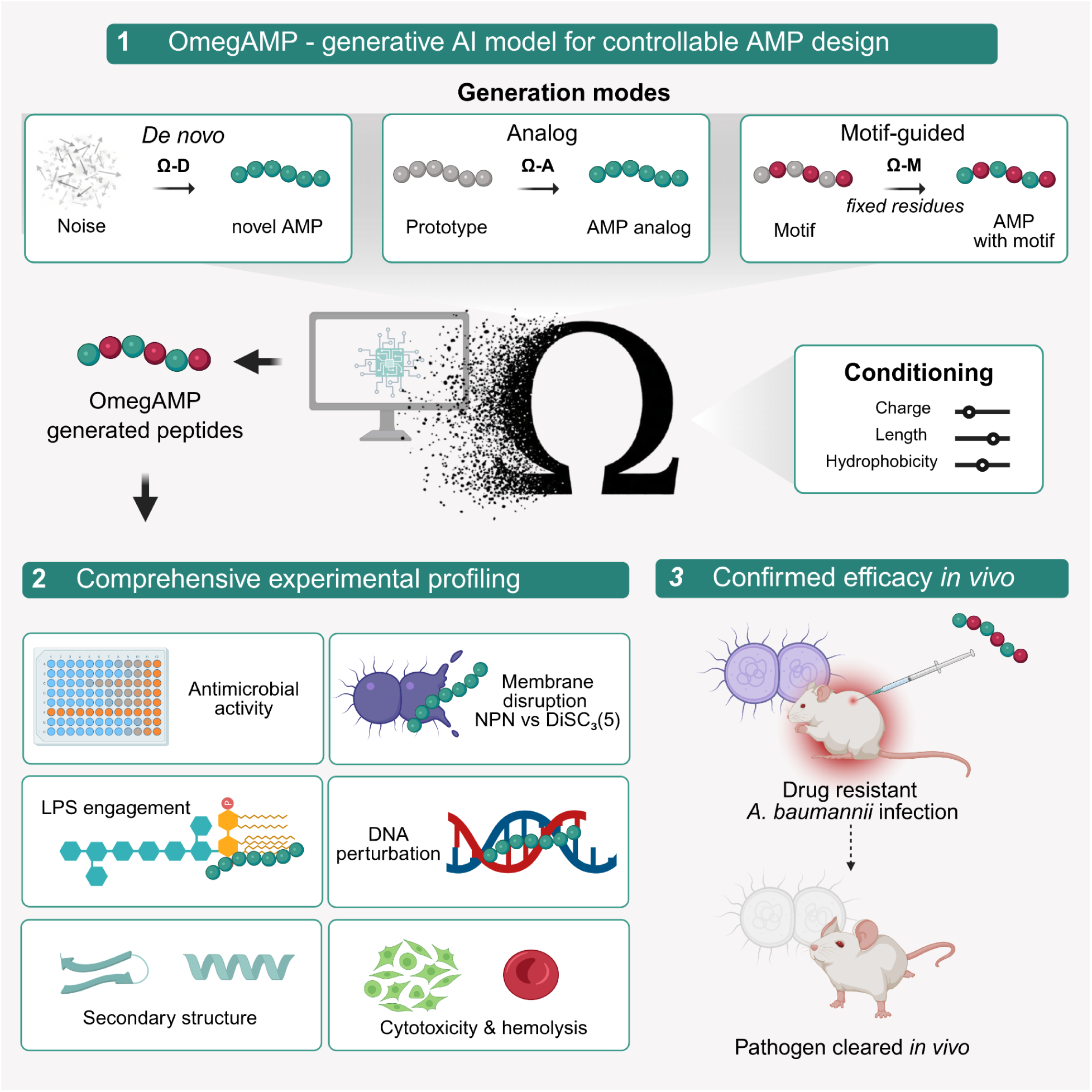

## Main

Antimicrobial resistance (AMR) is a leading and growing cause of global mortality^1^, with rising prevalence in serious hospital-acquired infections caused by carbapenem-resistant Gram-negative bacteria and methicillin-resistant *Staphylococcus aureus* (MRSA)^2^. Few drug candidates in the antibacterial pipeline address World Health Organization (WHO) critical-priority pathogens through mechanisms distinct from existing antibiotic classes, with limited approved options against carbapenem-resistant *Acinetobacter baumannii* (CRAB)^3,4^. New antimicrobials acting on conserved bacterial features rather than on specific mutable protein targets are therefore needed.

Antimicrobial peptides (AMPs) are a candidate class addressing this need. AMP bactericidal activity arises directly from a physicochemical interaction with the bacterial membrane, which enables bilayer insertion and disruption, leading to leakage of cellular contents and bacterial death^5^. Because the membrane lipid composition that AMPs exploit is broadly conserved and cannot be remodeled without fitness cost, AMPs select for resistance more slowly than conventional antibiotics^6^. Three key sequence-derived physicochemical properties shape AMP activity^5,7^. *Net charge*, typically +3 to +10 in active AMPs, drives the initial electrostatic attachment to the anionic bacterial surface and establishes selectivity against zwitterionic mammalian membranes. *Mean hydrophobicity*, typically between –0.2 and +0.5 on the Eisenberg scale, determines how readily the peptide inserts into the lipid bilayer and serves as a computable proxy for amphipathic architecture, which cannot be inferred from sequence alone. *Sequence length*, typically 15–50 residues, constrains both secondary structure formation and synthesis feasibility. Beyond these global properties, many AMPs carry short recurrent sequence motifs that confer auxiliary functions, expanding the antimicrobial mode of action beyond direct membrane disruption. Motifs that bind the lipid A moiety of bacterial lipopolysaccharide (LPS) sequester this primary endotoxin^8,9^, while nucleic acid binding motifs engage intracellular targets such as DNA^9,10^.

Generative AI models are increasingly applied to AMP design^11–18^. However, existing models either condition on coarse activity labels or rely on *post-hoc* filtering, neither of which gives precise control over the physicochemical properties that determine AMP function^15,16^. This is also the case for the analog generation task, introduced for AMPs by HydrAMP^11^, as existing tools cannot constrain the analog’s physicochemical profile relative to its prototype. Finally, no current AMP model fixes specific residues for functional motifs while optimizing the surrounding sequence, an approach validated in protein design through motif-fixing diffusion^19,20^, that would enable the design of AMPs retaining auxiliary functions such as LPS engagement or DNA perturbation.

We reasoned that the two additional generation modes could offer control beyond the physicochemical properties we condition on directly. Because analogs by definition retain sequence identity with their prototype, we hypothesized that they inherit properties we never specify nor optimize directly, such as membrane-disruption mechanism and safety, so that prototype choice becomes a design lever in its own right. Motif-guided generation raised an even sharper question, whether antimicrobial activity could be installed onto a non-antimicrobial scaffold while preserving its original function, such as LPS engagement or DNA perturbation.

We present OmegAMP, a conditional diffusion model for AMP design with explicit control over net charge, mean hydrophobicity, and sequence length. A single framework supports *de novo* design under physicochemical property targets, analog generation from existing peptide prototypes, and motif-guided design that preserves user-specified functional residues. We synthesized and wet-lab characterized OmegAMP-designed peptides covering all three modes alongside their reference prototypes, measuring antimicrobial activity against a panel of bacterial strains including multidrug-resistant (MDR) isolates, mammalian cell cytotoxicity and hemolysis, mode of bacterial-membrane disruption, secondary structure, and, where relevant, LPS engagement or DNA perturbation. *De novo* generation yielded potent antimicrobials with broad activity against MDR Gram-negative isolates. Analog generation converted inactive prototypes into potent antimicrobials, with prototype choice shaping the analogs’ mode of bacterial-membrane disruption and, through it, their mammalian-cell safety profile. Motif-guided generation preserved user-specified functional residues across modes. For AMP prototypes that engage LPS, analogs matched or exceeded prototype LPS engagement while remaining antimicrobial. A redesign of a non-antimicrobial leucine zipper acquired antimicrobial activity and exceeded the prototype’s DNA perturbation. Selected leads showed efficacy in murine skin scarification and thigh infection models. OmegAMP establishes controllable conditional generation as a general-purpose tool for antimicrobial peptide design, where the choice of prototype shapes not only sequence properties but the mechanism and safety profile of the resulting peptides.

## Results

### Physicochemistry-based conditional diffusion framework enables controlled peptide generation and motif preservation

OmegAMP is organized around a separation of physicochemical conditioning from the generation modes (**Figure 1**). During training, peptide embeddings derived from amino acid sequences are noised to varying degrees, and the model learns to denoise them conditioned on the corresponding conditioning vector **(Figure 1A)**. At inference, given a conditioning vector, the model iteratively denoises an initial noisy embedding toward a valid peptide representation which is then decoded into an amino acid sequence (**Figure 1B**). *De novo generation* (denoted OmegAMP-D) begins from pure noise, exploring the full space of peptide sequences consistent with the conditioning vector. Two additional generation modes introduce constraints on the sequence, by modifying the diffusion trajectory (**Figure 1C**). *Analog generation* (denoted OmegAMP-A) is initialized from a partially noised embedding derived from a prototype peptide, where *the exploration strength* (*τ*) controls how much noise is added, balancing fidelity to the prototype sequence against exploration toward novel peptides. In *motif-guided generation* (OmegAMP-M), designated residues are held fixed throughout denoising, optimizing the surrounding sequence while preserving a provided motif. Analog and motif-guided modes can operate jointly, enabling exploration with respect to a prototype sequence while simultaneously preserving a predefined motif.

**Figure 1.**
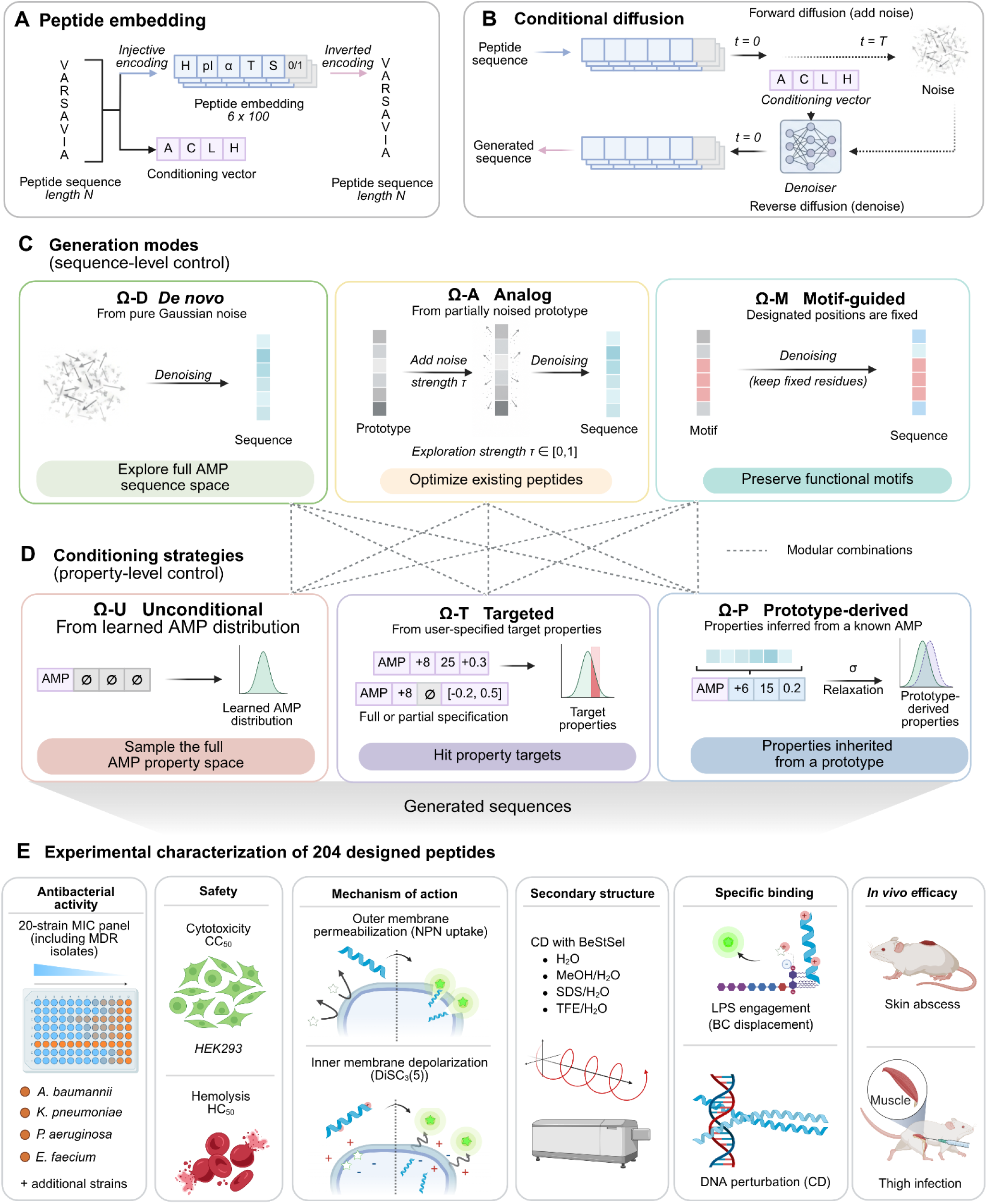
OmegAMP integrates wet lab-validated property control, prototype-guided optimization, and motif preservation. **(A)** Peptide embedding. Each residue maps to a 6D vector encoding hydrophobicity (H), isoelectric point (pI), α-helix propensity (α), transmembrane tendency (T), selectivity (S), and a padding token (0/1). A separate conditioning vector encodes target antimicrobial activity (A), charge (C), length (L), and mean hydrophobicity (H). **(B)** Conditional diffusion. During training, peptide embeddings are progressively noised over t = 0 → T, and a denoiser learns to reverse the process conditioned on the conditioning vector. At inference, iterative denoising of an initial noisy embedding yields a sequence. **(C)** Generation modes. *De novo* (OmegAMP-D), generation from pure Gaussian noise; analog (OmegAMP-A), generation from a partially noised prototype with exploration strength τ ∈ [0, 1]; motif-guided (OmegAMP-M), generation in which designated residues are constrained to stay fixed throughout denoising. **(D)** Conditioning strategies. Unconditional (OmegAMP-U), sampling from the learned AMP distribution; targeted (OmegAMP-T), explicit values or ranges for activity, charge, length, or hydrophobicity; prototype-derived (OmegAMP-P), conditioning vector et to the prototype peptide’s net charge, length, and hydrophobicity, with relaxation σ ∈ [0, 1] controlling how closely the generation matches that target. Modes and strategies combine modularly (e.g., OmegAMP-DP, OmegAMP-DT, OmegAMP-AP, OmegAMP-AT, OmegAMP-AMT). **(E)** Experimental characterization. MIC across a 20-strain panel including 8 MDR pathogens, including *A. baumannii*, *K. pneumoniae*, *P. aeruginosa*, *E. faecium*; HEK293T cytotoxicity (CC_50_) and hemotoxicity (HC_50_); outer– and cytoplasmic-membrane disruption (NPN, DiSC3(5)); secondary structure by circular dichroism (CD) in H_2_O, MeOH/H_2_O, SDS/H_2_O, and TFE/H_2_O; LPS engagement by BODIPY-cadaverine displacement; DNA perturbation by CD; and murine skin scarification and thigh infection.

To obtain control over physicochemical properties, we expose the model during training to conditioning vectors that are derived from the underlying peptides and are subsequently randomly masked. This training schema yields conditioning vectors that span from fully defined to fully masked across charge, length, and hydrophobicity, while the antimicrobial activity indicator remains set to whether the peptide is an AMP/non-AMP. The model thereby learns associations between peptide embeddings and their target properties, enabling flexible property control at inference. We propose three complementary strategies to populate the conditioning vector with target profiles (**Figure 1D**). For *unconditional generation*, the conditioning vector is left entirely unspecified except for the prerequisite of antimicrobial activity (OmegAMP-U), allowing the model to sample from the learned distribution of all training AMPs. For *targeted conditioning* (OmegAMP-T), the conditioning vector is either partially or fully populated with explicit values or ranges for any combination of charge, length, hydrophobicity, and antimicrobial activity. For *prototype-derived* conditioning (OmegAMP-P), target profiles are derived from prototype peptides, with a *relaxation* parameter (*σ*) controlling how closely the generated peptides should match the prototypes’ physicochemical profiles.

Generation and conditioning strategies can be used in combination, offering flexibility which enables practitioners to tailor the generative process to specific design objectives: preserving a functional motif while optimizing for antimicrobial activity, exploring structural variants of a lead peptide within defined hydrophobicity bounds, or systematically varying charge while maintaining sequence similarity to a validated AMP. We use a modular nomenclature to denote the specific combination of generation modes and conditioning strategies. For analog generation, OmegAMP-AT uses targeted conditioning and OmegAMP-AP derives the target property profile from the prototype. For motif-guided design, OmegAMP-MT simultaneously preserves key functional residues and optimizes specific physicochemical properties.

The experimental campaign tested 215 peptides across the three generation modes (**Figure 1E**): 95 *de novo* peptides (Ω-DP, Ω-DT), 59 analogs (Ω-AP, Ω-AT, Ω-AU) of 6 inactive prototypes, and 50 motif-preserving designs of LPS-engaging (Ω-AMT) or DNA-perturbing (Ω-MT) prototypes, alongside 11 prototypes and the antibiotics polymyxin B and levofloxacin as references. All peptides were characterized across antimicrobial activity, cytotoxicity, hemotoxicity, mechanism of action, and secondary structure (**Figure 1E**). From these results we selected 11 leads representing each generation mode, based on strain coverage (number of strains reached at MIC ≤ 2 μM), overall potency (MIC₅₀, the median MIC across the 20-strain panel), and safety (hemolysis HC₅₀ and cytotoxicity CC₅₀) (**Table 1**). These leads were advanced to murine *in vivo* characterization in skin scarification and thigh infection models.

**Table 1.**
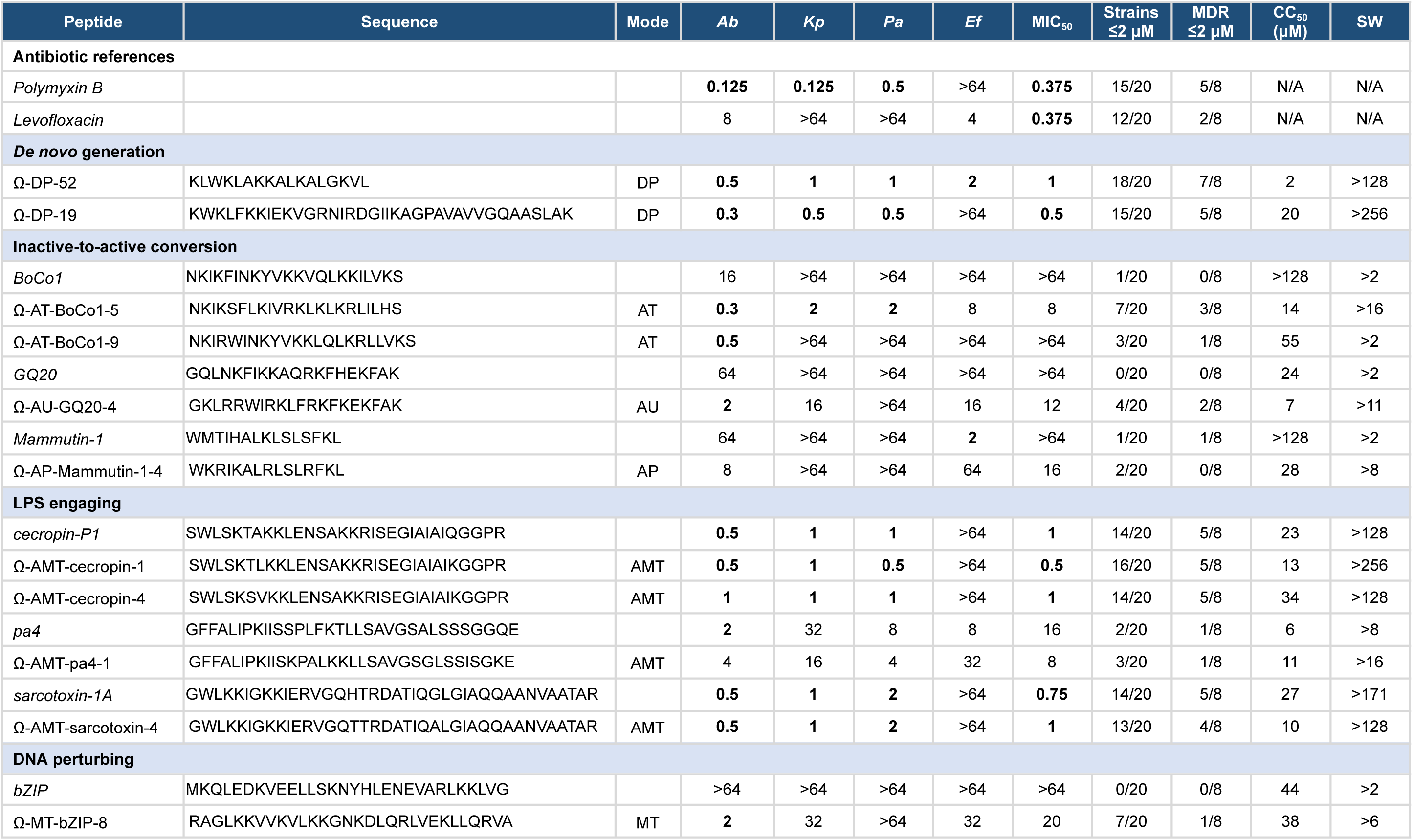
Selected OmegAMP leads across generation modes. Strain abbreviations: Ab, *A. baumannii* ATCC BAA-1605 (carbapenem-resistant); Kp, *K. pneumoniae* ATCC BAA-2342 (carbapenem-resistant); Pa, *P. aeruginosa* ATCC BAA-3197 (fluoroquinolone-resistant); Ef, *E. faecium* ATCC 700221 (vancomycin-resistant). MIC values are in μM. Bold indicates MIC ≤ 2 μM. Strains ≤ 2 μM reports count across the 20-strain panel, and MDR ≤ 2 μM count across eight MDR isolates. SW, safety window (HC₅₀/MIC₅₀). Prototype rows are italicized. Polymyxin B and levofloxacin are reference antibiotics.

### Conditioning achieves precise property control within biologically accessible boundaries

To contextualize OmegAMP’s conditioning against the experimentally observed AMP design space, we analyzed property-activity relationships across peptides from a curated database^21^ and by predicting the activity from the properties with various machine learning models. Potent activity concentrated at higher positive net charge, slightly negative to near-to-zero mean hydrophobicity, and lengths of 15–25 amino acid residues (**Figure 2A**; **Extended Data Fig. 1A-C**). Aggregation propensity rose with hydrophobicity and fell with net charge (**Figure 2B**; **Extended Data Fig. 1D-F**). Across these peptides, net charge was the strongest determinant of activity and remained dominant after controlling for mean hydrophobicity and length (**Figure 2C**). Hydrophobicity contributed weakly and largely indirectly, acting through amphipathic architecture and aggregation rather than as a direct determinant. These properties together only partially predicted MIC. In cross-validation, predictive performance improved from linear to random forest models and was highest when amino acid composition was added to the physicochemical descriptors, indicating that potency depends on sequence features beyond net charge, mean hydrophobicity, and length (**Figure 2D**). Since charge and hydrophobicity were anti-correlated, peptides combining high charge with high hydrophobicity occupied a narrow, aggregation-prone region.

**Figure 2.**
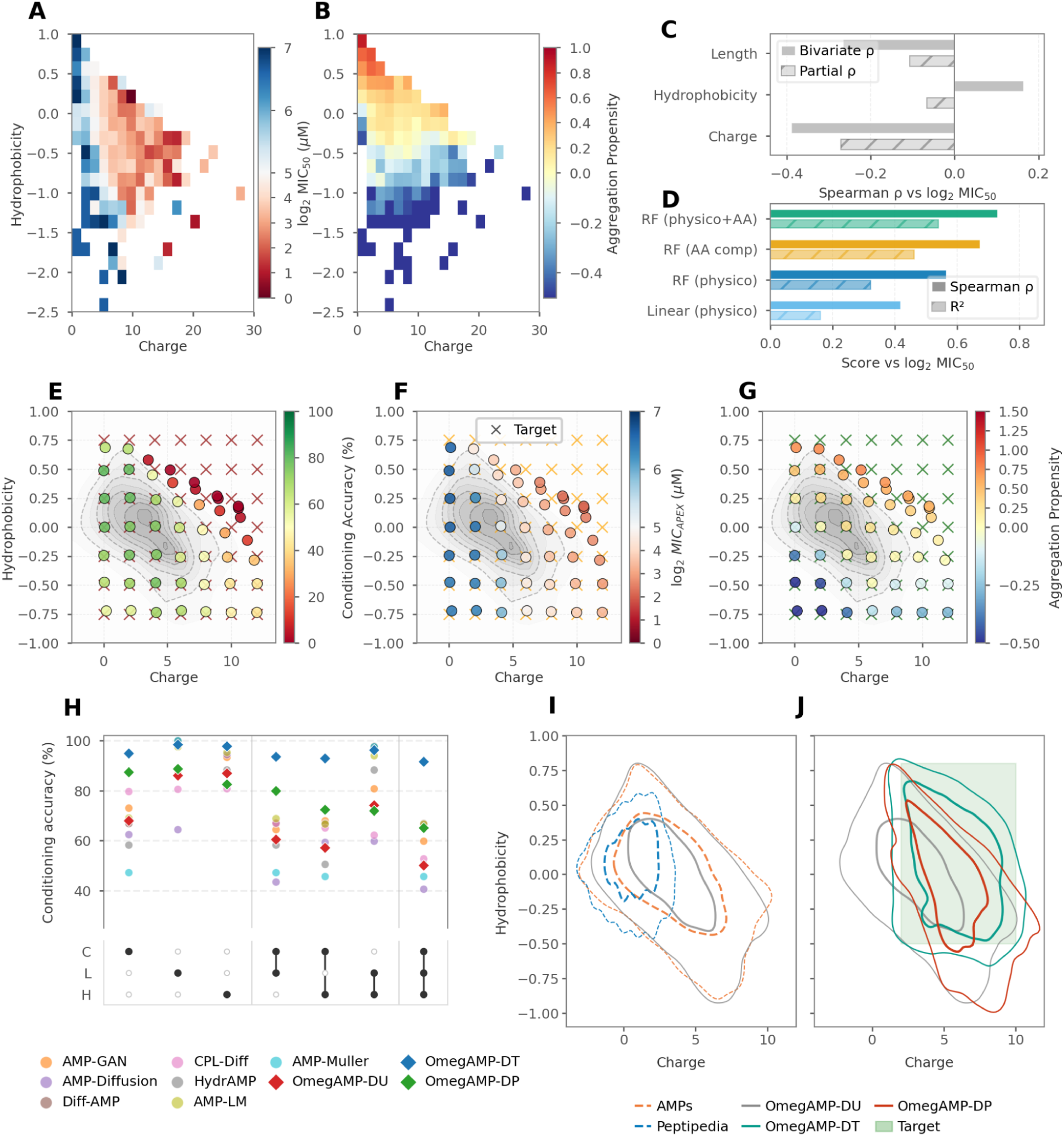
OmegAMP maps and controls the feasible physicochemical design space. (**A**) MIC across charge-hydrophobicity space (median across available species, log₂ MIC₅₀, μM); *n = 5,207* AMP-database peptides (see also **Extended Data Fig. 1A to 1C**). Red marks lower MIC. White cells indicate charge-hydrophobicity bins containing no peptides. (**B**) Aggrescan aggregation propensity across the same space and peptides as (A) (see also **Extended Data Fig. 1D to 1F**). Red marks higher aggregation. (**C**) Correlation of charge, hydrophobicity, and length with log₂ MIC₅₀ for the peptides in (A) and (B) (bivariate Spearman ρ, solid; partial ρ controlling for the other two properties, hatched). (**D**) Cross-validated log₂ MIC₅₀ prediction by linear regression and random forest (RF) models trained on physicochemical descriptors (“physico” = charge, hydrophobicity, length), amino acid composition (“AA comp”), or both (Spearman ρ, solid; R², hatched). (**E**) Conditioning accuracy across the charge-hydrophobicity grid (49 coordinates, 500 peptides per coordinate), the fraction of peptides within ±0.25 charge units, ±0.1 hydrophobicity units, and exact length match (see also **Extended Data Fig. 1G to 1L**). Gray contours indicate training data density; crosses denote target coordinates. (**F**) Predicted activity (log₂ MIC_APEX_, μM) across the grid in (E) (see also **Extended Data Fig. 1M to 1O**). Conventions as in (E). (**G**) Predicted aggregation propensity across the grid in (E) (see also **Extended Data Fig. 1P to 1R**). Conventions as in (E). (**H**) Benchmarking conditioning accuracy against published AMP generative models under single-property, pairwise, and three-property constraints (charge 2-10, length 5-30, hydrophobicity –0.5 to +0.8). Filled circles, baseline methods; diamonds, OmegAMP-DU (red), OmegAMP-DT (blue), and OmegAMP-DP (green). (**I**) Property-space coverage of unconditional generation (gray solid; *n = 50,000*) overlaid on curated AMPs (orange dashed; *n = 35,878*) and general peptides from Peptipedia (blue dashed; *n = 774,405*). (**J**) Property-space coverage of OmegAMP-DU (gray), OmegAMP-DT (teal), and OmegAMP-DP (red) generation (*n = 50,000* each) relative to the target range (charge 2-10, length 5-30 residues, and hydrophobicity **-**0.5 to +0.8; green shading).

While OmegAMP achieves high-fidelity single property control in targeted conditioning, multiproperty control results in deviations at simultaneously high charge and hydrophobicity targets, reflecting intrinsic physicochemical constraints (**Extended Data Fig. 1G-I**). Cationic residues inherently reduce mean hydrophobicity, which is a fundamental limitation of amino acid composition. To systematically map the controllable property space, we generated peptides across a grid of 49 target combinations for each property pair: net charge (0 to +12 in increments of 2), mean hydrophobicity (–0.75 to +0.75 in increments of 0.25), and length (5–35 amino acid residues in increments of 5). Across all property grids, conditioning accuracy was quantified as the fraction of generated peptides achieving net charge within ±0.25 units, mean hydrophobicity within ±0.1 units, and exact length match. These tolerances fall below the effect of a single residue substitution, corresponding to stringent criteria for conditioning success. The model achieves high conditioning accuracy (>80%) across most of the net charge-mean hydrophobicity space, yet with reduced performance in the high-net charge, high-mean hydrophobicity region (**Figure 2E; Extended Data Fig. 1J**), consistent with the empirically observed compositional constraint described above.

The same challenging region resides over the edge of the region of the highest predicted antimicrobial activity (**Figure 2F**). Peptides with net charge +6 to +10 and mean hydrophobicity +0.25 to +0.5 show the lowest mean predicted MIC across bacterial strains from a pretrained activity predictor APEX^22^ (MIC_APEX_). This region also exhibits elevated aggregation propensity (**Figure 2G**), recapitulating the experimental landscape (**Figure 2B**). When targets become compositionally infeasible (e.g., net charge +12 for a sequence of 8 residues), the model generates peptides at the boundary of achievable property space (**Extended Data Fig. 1K**). Mean hydrophobicity-length combinations showed no such limitation (**Extended Data Fig. 1L**), likely because mean hydrophobicity averages over all residues, offering greater compositional flexibility than net charge, which depends on discrete cationic residues. Predicted activity patterns across these dimensions favored shorter sequences (5–25 residues) and elevated net charge, consistent with experimental observations (**Extended Data Fig. 1M-O** cf. **Extended Data Fig. 1A-C**), while predicted aggregation propensity rose with hydrophobicity across the same dimensions (**Extended Data Fig. 1P-R** cf. **Extended Data Fig. 1D-F**). Collectively, these patterns demonstrate that targeted conditioning successfully generates sequences for compositionally plausible targets, while defaulting to the boundary of achievable property space when targets become infeasible.

Next, we evaluated conditioning accuracy for generating peptides not for specific target property values, but within property ranges: net charge (2–10), length (5–30 residues), and mean hydrophobicity (–0.5 to +0.8), encompassing the therapeutically relevant AMP property space. In this task, we compared OmegAMP to seven published AMP generative models across single-property, pairwise, and multi-property constraints (**Figure 2H**). Conditioning accuracy is defined as the fraction of generated peptides whose properties fall within the specified target range. For baselines lacking explicit property control (AMP-GAN, AMP-Diffusion, Diff-AMP, AMP-Muller, AMP-LM, HydrAMP, CPL-Diff), the metric scores their default sampling distribution against the target range. OmegAMP with targeted conditioning (OmegAMP-DT) achieved >90% success across all property combinations, including the most demanding scenario requiring simultaneously meeting all three constraints (C+L+H). OmegAMP without conditioning (OmegAMP-DU), which samples properties from the learned AMP distribution rather than explicit targets, achieved 70–80% accuracy. Alternative approaches showed variable performance on single-property tasks and generally degraded as constraints were combined, consistent with these models lacking a mechanism to enforce property targets at generation time.

This accuracy difference between conditioning strategies reflects their distinct coverage of property space (**Figure 2I, 2J**). When the conditioning vector is left unspecified (OmegAMP-DU), the model samples from the learned distribution of training AMPs, spanning a broad property space that includes sequences with low or negative charge and extreme hydrophobicity values (**Figure 2I**). When conditioning vectors are derived from prototype peptides with demonstrated antimicrobial activity (OmegAMP-DP), the resulting distribution shifts toward property combinations characteristic of functional AMPs (**Figure 2J**). When the conditioning vector is explicitly populated with target values (OmegAMP-DT), the model concentrates output within the specified boundaries. These distinct conditioning strategies offer complementary approaches to navigating the physicochemical space, with the choice depending on whether the design objective favors broad exploration, emulation of known active profiles, or precise property specification. This flexibility illustrates the practical value of the conditioning framework for therapeutic peptide design.

### Analog generation enables controlled optimization of existing peptide prototypes

*De novo* generation offers broad exploration of sequence space, yet many practical applications require the optimization of existing peptides. This scenario parallels lead optimization in drug discovery, where promising compounds undergo iterative refinement to improve potency or selectivity while preserving core structural features^23^. We previously demonstrated that analog generation can both improve inactive peptides and further optimize active ones^11^. OmegAMP extends analog generation through a two-parameter setup, which decouples sequence exploration strength (τ, the noise applied to the prototype) from property relaxation (σ, the deviation permitted from the prototype’s physicochemical profile). We evaluated analog generation *in silico* on 500 active and 500 inactive experimentally validated prototypes, generating 10 analogs per configuration across τ = 0.0–0.7 and σ = 0.0–1.0, with activity predicted by APEX and similarity measured by normalized local alignment (**Extended Data Fig. 2**, **Data S1**). Property deviation scaled approximately linearly with σ, while σ = 0 preserved prototype properties across all τ (**Extended Data Fig. 2A-C**). Adopting ≥60% sequence similarity as the threshold for analog identity, both prototype classes optimized best at σ = 1.0, with potency improving as τ increased, but they differed in how quickly similarity eroded. Active prototypes tolerated aggressive exploration, reaching a median MIC_APEX_ of 4.4 μM while still 64% similar at τ = 0.7 (**Extended Data Fig. 3A and 3B**). Inactive prototypes instead crossed the 60% floor already at τ = 0.3, capping their best in-threshold potency at 10.6 μM (61% similar) (**Extended Data Fig. 3C and 3D**). Optimizing inactive prototypes therefore trades more sequence identity for a smaller potency gain (**Extended Data Fig. 2E and 2F**, with detailed parameter sweeps in **Extended Data Fig. 3**). Across the five analog conditioning strategies (unconditional AU, property-targeted AT, prototype-derived AP, motif-guided AM, and motif-plus-property AMT), property-targeted OmegAMP-AT achieved the lowest median MIC_APEX_ and significantly outperformed HydrAMP-A, the state of the art for AMP optimization, on both prototype classes (**Extended Data Fig. 2G and 2H**; Mann-Whitney U, p < 0.001), while OmegAMP returned far more valid analogs (≥494/500 vs. as few as 94/500 for HydrAMP-A at *τ = 1,* **Extended Data Fig. 2G and 2H**) and explored sequence space more broadly (**Extended Data Fig. 2I and 2J**).

**Figure 3.**
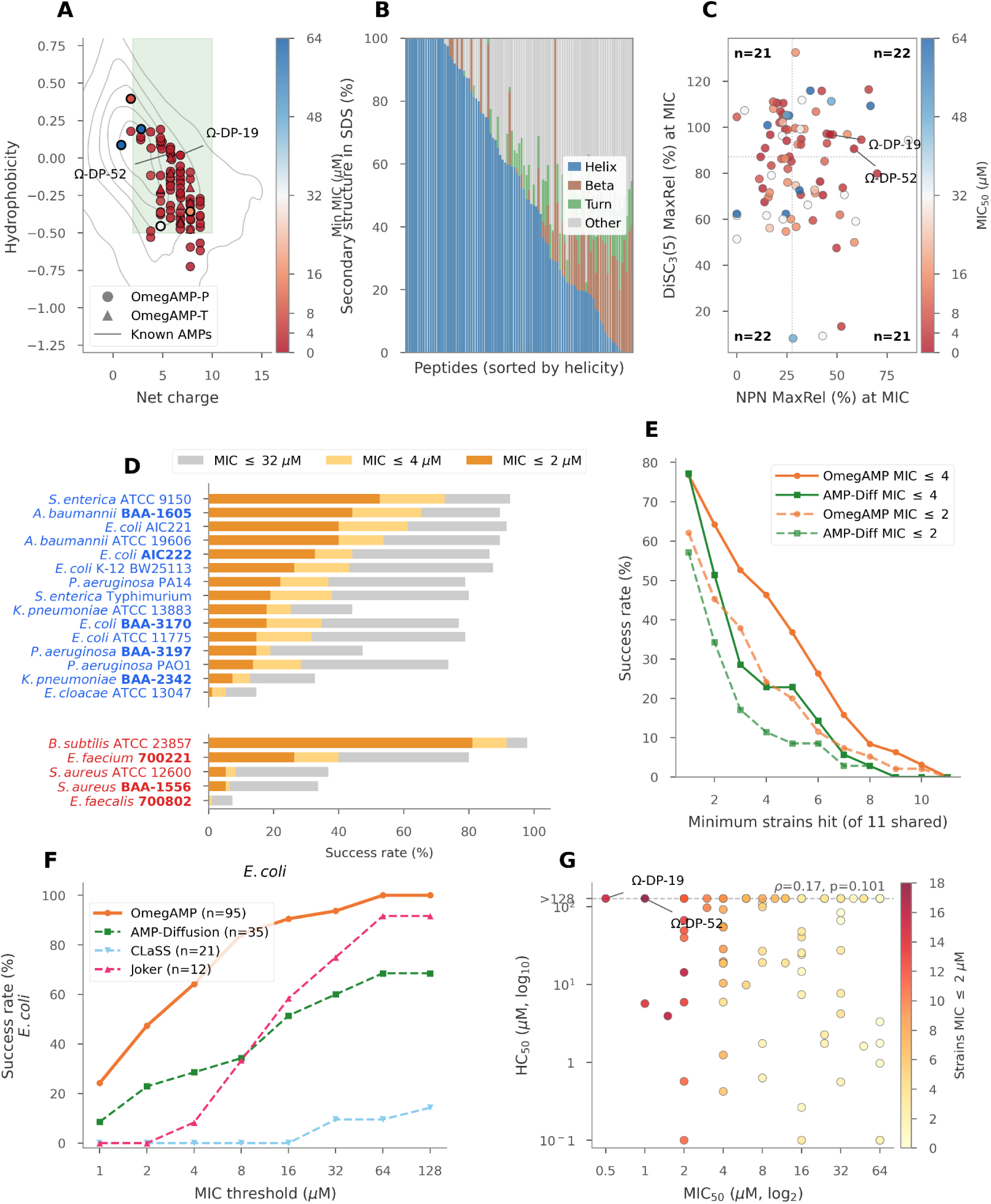
Controlled *de novo* generation yields potent and mechanistically diverse antimicrobials. (**A**) Physicochemical coverage of OmegAMP-DP (circles, *n = 85*) and OmegAMP-DT (triangles, *n* = 10) peptides relative to known AMPs (*n* = 35,878). Points colored by MIC₅₀ across the 20-strain panel; thicker outlines, inactive peptides. Gray contours, density of known AMPs; green shading, therapeutic target range (charge 2–10, hydrophobicity –0.5 to +0.8); Ω-DP-19 and Ω-DP-52 labeled. (**B**) Secondary structure composition in SDS micelles by CD spectroscopy with BeStSel deconvolution. Bars show fractional α-helix, β-strand, turn, and other content, sorted by decreasing helicity (see also **Extended Data Fig. 4A-C**). (**C**) Membrane disruption against *A. baumannii* ATCC 19606, each peptide assayed at its MIC. NPN uptake reports outer membrane permeabilization; DiSC₃(5) fluorescence reports cytoplasmic membrane depolarization. Points colored by MIC₅₀. Dashed lines, medians, partitioning peptides into NPN-dominant, DiSC-dominant, both-strong, and both-weak classes; *n* per quadrant indicated. Ω-DP-19 and Ω-DP-52 labeled. (**D**) Activity across the 20-strain panel by Gram stain. Bars show the fraction of *de novo* peptides reaching MIC ≤ 2 μM (dark orange), ≤ 4 μM (light orange), or ≤ 32 μM (gray). MDR isolates in bold; strain labels in blue (Gram-negative) or red (Gram-positive). (**E**) Cross-study success rates across 11 strains also reported for AMP-Diffusion (*n = 35*). Curves show the fraction of peptides with MIC ≤ 4 μM (solid) or MIC ≤ 2 μM (dashed) against at least the indicated minimum number of strains (x-axis). (**F**) Benchmarking against published generative models on *E. coli* strains: OmegAMP (*n* = 95), AMP-Diffusion (*n* = 35), CLaSS (*n* = 21), Joker (*n* = 12). μg/mL values converted to μM. (**G**) Hemolysis (HC₅₀) and potency (MIC₅₀) for *de novo* peptides; points colored by the number of strains against which the peptide reached MIC ≤ 2 μM; HC₅₀ right-censored at 128 μM. Ω-DP-19 and Ω-DP-52 labeled. Spearman ρ and *p* annotated (see also **Extended Data Fig. 4D,E**).

### Conditioned *de novo* generation yields potent and mechanistically diverse antimicrobials

Having established control *in silico*, we tested whether specified physicochemical profiles translate into antibacterial function. We characterized 215 peptides *in vitro*, beginning with *de novo* generation. To evaluate whether precise physicochemical control translates to functional antimicrobial activity, we synthesized and tested 95 peptides generated *de novo* using two conditioning strategies: targeted conditioning (OmegAMP-DT, *n = 10*), where property values were explicitly specified, and prototype-derived conditioning (OmegAMP-DP, *n = 85*), where each designed peptide’s conditioning vector (C, H, L) was taken from an active AMP in DBAASP^21^ (MIC ≤ 32 μg/mL against at least one strain; see Methods), with a different prototype drawn per each design. Peptides were tested against a panel of 20 bacterial pathogens, including 15 Gram-negative and 5 Gram-positive strains.

The selected peptides spanned the therapeutically relevant property space with net charge ranging from +1 to +9 and mean hydrophobicity from –0.73 to +0.40 (**Figure 3A**). Of 95 peptides tested, 90 (95%) exhibited MIC ≤ 4 μM against at least one strain and 82 (86%) reached MIC ≤ 2 μM. The five inactive designs fell into two groups: three carried lower net charge (+1 to +3), consistent with the requirement for cationicity in membrane targeting, while the remaining two carried typical net charge but markedly negative mean hydrophobicity, indicating that amphipathicity, not net charge alone, is limiting.

Secondary structure composition was determined by CD spectroscopy with BeStSel^24^ deconvolution in SDS micelles, a membrane-mimetic environment (**Figure 3B**). Most peptides were unstructured in water (**Extended Data Fig. 4A**) but adopted ordered conformations in SDS, with helical folds dominating (≥ 50%); the remainder split between β-rich and disordered conformations. The *de novo* set thus spans the structural diversity of natural AMPs (α-helices, β-rich folds, disordered peptides that fold on membrane contact). Cross-medium CD spectra are reported in **Extended Data Fig. 4A-C**. Helical peptides reached lower MIC₅₀ than β-rich peptides (Mann-Whitney, MW *p < 0.001,* **Extended Data Fig. 4D**), with the helical-disordered difference not significant. Sequence novelty was heterogeneous across the *de novo* set (**Extended Data Fig. 4E**), but antimicrobial potency was uncorrelated with training identity (*p* > 0.05), indicating that the model produces functional peptides across the novelty spectrum.

Mechanistic profiling further showed that *de novo* generation did not collapse onto a single membrane phenotype. At each peptide’s MIC against *A. baumannii* ATCC 19606, NPN uptake measured outer-membrane permeabilization and DiSC₃(5) fluorescence measured cytoplasmic-membrane depolarization (**Figure 3C**). Peptides populated all four median-defined classes (i.e., NPN-dominant, DiSC-dominant, both strong, and both weak) revealing mechanistic heterogeneity within a library generated from the same global property controls.

Activity was preferentially directed against Gram-negative strains (**Figure 3D**), consistent with cationic AMPs engaging the anionic outer membrane more readily than the thicker peptidoglycan and modified lipid surfaces of Gram-positive envelopes^25^. All Gram-negative species were reached at MIC ≤ 2 μM, with *A. baumannii* BAA-1605 the most accessible MDR isolate (44%) and *K. pneumoniae* BAA-2342 the least (7%). Among Gram-positives, only *B. subtilis* ATCC 23857 (81%) and *E. faecium* 700221 (26%) were broadly susceptible at MIC ≤ 2 μM; the remaining Gram-positive strains were largely resistant.

To place these hit rates in context, we compared them with published AMP-generation studies reporting wet-lab testing^17,18,26^. Across 11 strains shared with AMP-Diffusion, OmegAMP produced a larger fraction of peptides active at 2 μM and 4 μM (**Figure 3E**). Remaining methods synthesized 12–35 peptides each; on *E. coli* at MIC ≤ 2 μM, OmegAMP produced 45 hits out of 95 candidates, versus 8 out of 35 for AMP-Diffusion and none out of 12 (Joker) or 21 (CLaSS) candidates (**Figure 3F**).

Across the *de novo* set, potency and safety varied independently (Spearman *ρ = 0.17, p = 0.105*, **Figure 3G**). Two peptides occupied the high-HC₅₀, low-MIC₅₀ corner and were selected to advance to murine experiments (**Table 1**). Ω-DP-19 had the highest safety window in the *de novo* set, with no detectable hemolysis (HC₅₀ >128 μM) and low mammalian cell cytotoxicity (HEK293 CC₅₀ = 19.8 μM), reaching MIC ≤ 2 μM against all five drug-resistant Gram-negatives. Ω-DP-52 achieved the broadest spectrum (MIC ≤ 2 μM against 18 of 20 strains, including MRSA and vancomycin-resistant *E. faecium* 700221), with no detectable hemolysis (HC₅₀ >128 μM) but higher cytotoxicity (CC₅₀ = 2.0 μM).

### Analog generation converts inactive peptides into active antimicrobials with prototype-determined safety profiles

We next asked whether the same framework could recover antibacterial activity from experimentally inactive sequences. To this end, we characterized the resulting analogs for activity, safety, and the membrane-disruption mechanism. We selected six prototype peptides experimentally characterized as inactive (MIC ≥ 128 μg/mL aross all reported strains in DBAASP): As-CATH4-6L^27^, Mammutin-1^22^, BoCo1^28^, GQ20^29^, DeNo1047^30^, and OP-145-TII4^11^. Notably, OP-145-TII4 was itself generated by HydrAMP as an analog of OP-145, yet failed to achieve the intended activity, underscoring this task’s difficulty. We generated 59 analogs in total, 10 per prototype except 9 for As-CATH4-6L. These were distributed across three conditioning strategies: prototype-derived (OmegAMP-AP, *n = 22*), targeted (OmegAMP-AT, *n = 29*), and unconditional (OmegAMP-AU, *n = 8*). All analogs maintained ≥60% sequence similarity to their prototype. Analog sequence families are shown aligned to their prototypes in **Extended Data Fig. 5A**. Full MIC, HC_50_, and CC_50_ profiles for all 59 analogs are shown in **Extended Data Fig. 5B**.

Per-family hit rates at MIC ≤ 2 μM ranged from 5 (OP-145-TII4) to 10 (GQ20 and BoCo1) out of 10, with 50 of 59 analogs reaching this threshold against at least one strain across the six families (**Figure 4A**). As in the *de novo* set, the analogs were more potent against Gram-negative than Gram-positive strains (**Figure 4B**). *A. baumannii* BAA-1605 was the most accessible MDR isolate at MIC ≤ 2 μM, with the remaining Gram-negative MDR isolates reaching lower fractions and Gram-positive MDR isolates almost entirely resistant. To benchmark analog generation against a prior method, we compared OmegAMP-A and HydrAMP-A analogs on the four species both were assayed against: *E. coli*, *S. aureus*, *A. baumannii* and *P. aeruginosa* (**Extended Data Fig. 5C**). An analog counted as successful when it inhibited at least one of these species. OmegAMP-A achieved a higher success rate than HydrAMP-A at every threshold, 59.3% versus 19.2% of analogs at MIC ≤ 2 µM, 69.5% versus 30.8% at MIC ≤ 4 µM and 94.9% versus 84.6% at MIC ≤ 32 µM. The same ordering held on A. baumannii ATCC BAA-1605, the one strain targeted experimentally by both models, where the rates were 49.2% versus 15.4%, 54.2% versus 23.1% and 89.8% versus 80.8%.

**Figure 4.**
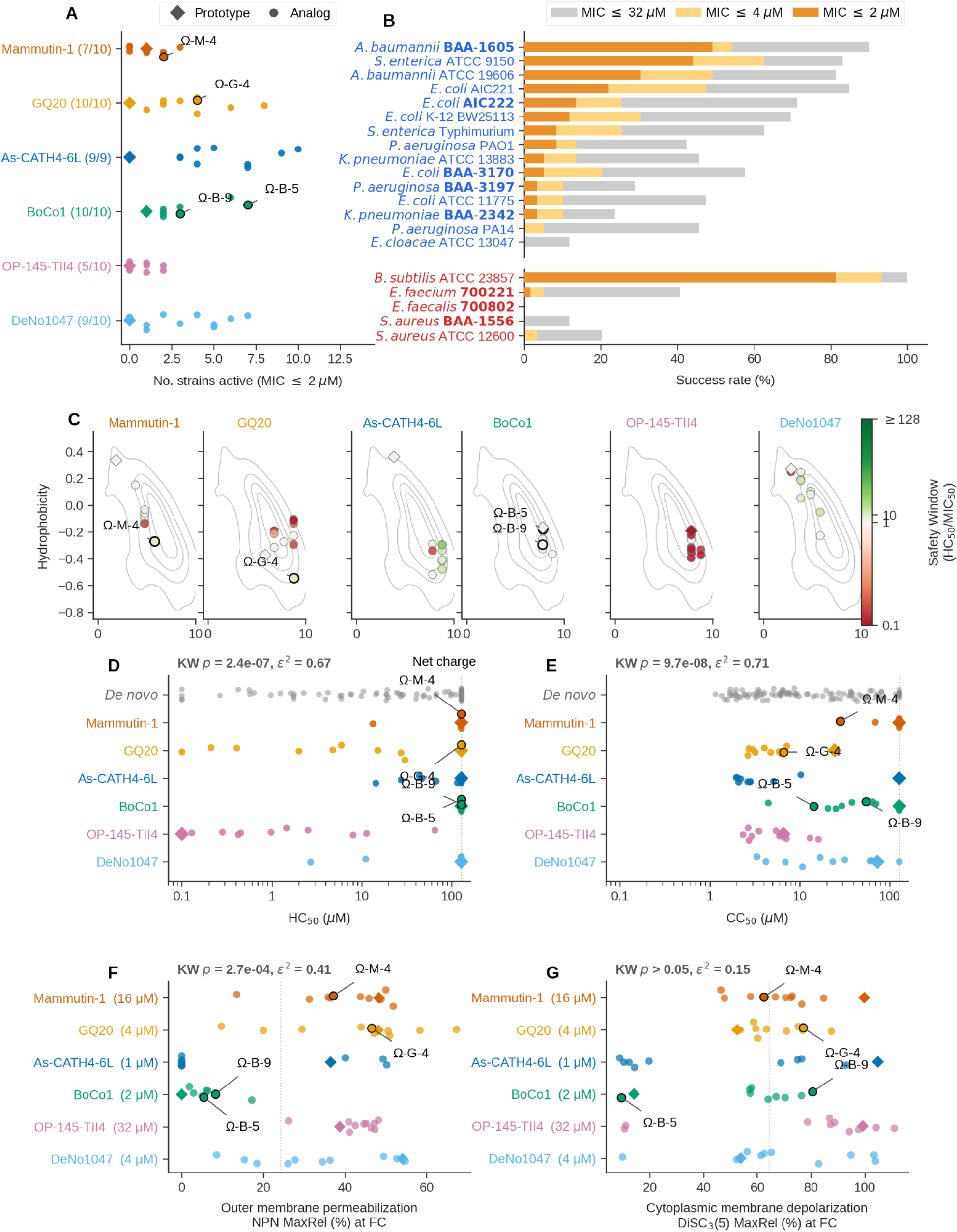
Analog generation converts inactive peptides into antimicrobials with prototype-determined mechanism and safety. (**A**) Number of strains with MIC ≤ 2 μM for analogs of six inactive prototypes. Diamonds, prototypes; fractions, analogs per family active at MIC ≤ 2 μM against at least one strain (see also **Extended Data Fig. 5A to 5C**). (**B**) Fraction of analogs reaching MIC ≤ 32, 4, and 2 μM per strain (Gram-negative, top; Gram-positive, bottom). MDR isolates in bold. (**C**) Net charge versus mean hydrophobicity by prototype family, colored by safety window (HC50/MIC50). Gray contours, property distribution of known active AMPs; diamonds, prototypes; points, analogs (see also **Extended Data Fig. 5D**). (**D**) HC₅₀ (μM) by prototype family; de novo peptides as reference (top row). Diamonds, prototypes; leads outlined in black. Shaded region at ≥128 μM, right-censored values. KW p and ε² report variance attributable to the prototype family. (**E**) CC₅₀ (μM) by prototype family. Conventions as in (D), with right-censoring at ≥128 μM. (**F**) Outer membrane permeabilization (NPN MaxRel, %) by prototype family, each family assayed at a single fixed concentration selected for that family and shown next to the family name (axis label FC). Diamonds, prototypes; leads outlined in black. KW p and ε² as in (D). (**G**) Cytoplasmic membrane depolarization (DiSC₃(5) MaxRel, %) by prototype family (see also **Extended Data Fig. 5E and 5F**). Conventions as in (F).

To understand how OmegAMP reached active sequences from inactive prototypes, we examined the analogs’ position in net charge-mean hydrophobicity space (**Figure 4C**) and their secondary structure fractions (**Extended Data Fig. 5D**). Three routes to activation emerged. The first, property-driven optimization, shifted analogs substantially in net charge-mean hydrophobicity space. Mammutin-1 analogs gained net charge and lost mean hydrophobicity (median ΔC +3, Δ<H> –0.4) but most remained disordered, while GQ20 analogs gained both net charge and mean hydrophobicity (ΔC +3, Δ<H> +0.15) and retained the prototype’s helical tendency. The second, property-preserving optimization, kept prototype properties nearly intact (|ΔC| ≤ 1, |Δ<H>| ≤ 0.15) while compositional or positional changes drove potency gains. OP-145-TII4 and BoCo1 analogs followed this route and retained their prototypes’ structural tendencies. The third, structural induction, was observed only in As-CATH4-6L, where the disordered prototype gave rise to nearly fully helical analogs. OP-145-TII4 was the peptide HydrAMP had previously failed to activate as an OP-145 analog. With OmegAMP it followed the property-preserving route rather than a disorder-to-helix transition, recovering activity in 5 of its 10 analogs. These analogs nonetheless retained the hemolytic and cytotoxic liabilities of their lineage, with HC₅₀ below 1.5 μM in most cases. DeNo1047 sat near the property-preserving boundary on net charge (ΔC +1) and mean hydrophobicity (Δ<H> –0.13) but its analogs spanned the full secondary structure range, combining property preservation with partial structural induction. Across the six families, helicity differed strongly by prototype (KW *p* < 0.001, ε² = 0.59; **Extended Data Fig. 5D**), reflecting the structural diversity that underlies the three routes.

Although OmegAMP optimizes for antimicrobial activity and does not condition on toxicity during generation, the resulting analogs acquire safety-relevant properties from their prototypes, so the choice of prototype offers a route to controlling mechanism and safety. KW tests across the six families rejected a common distribution for both HC₅₀ (*p* < 0.001, ε² = 0.64) and CC₅₀ (*p* < 0.001, ε² = 0.68), indicating that prototype family accounts for the majority of variance in analog safety (**Figure 4D, E**). Mammutin-1, BoCo1, and DeNo1047 produced largely non-hemolytic analogs (HC₅₀ near or above the assay ceiling), with cytotoxicity ranging from low (Mammutin-1) to moderate (BoCo1, DeNo1047); As-CATH4-6L, GQ20, and OP-145-TII4 yielded markedly less safe analogs, either hemolytic (lower HC₅₀) or cytotoxic (lower CC₅₀) on at least one axis.

To ask whether prototype family also shapes the membrane mechanism of the analogs, we measured outer-membrane permeabilization (NPN uptake) and cytoplasmic-membrane depolarization [DiSC₃(5) fluorescence] for each analog. Because each analog has a different MIC, comparing prototype effects across families requires controlling for concentration; we therefore tested each family at a single concentration chosen for that family (**Methods**; values indicated next to each row in **Figure 4F, G**). NPN response stratified by prototype family (**Figure 4F**; KW *p* < 0.001, ε² = 0.36), while DiSC₃(5) response did not (**Figure 4G**; *p* > 0.05). When the same assays were instead run at each analog’s MIC (**Extended Data Fig. 5E**, **F**), both readouts appeared to stratify by family [NPN ε² = 0.43; DiSC₃(5) ε² = 0.46]. Within the ≥60% similarity floor imposed during analog generation, the prototype family thus accounts for an increasing share of analog variance from outer-membrane permeabilization (**Figure 4F**) through helicity (**Extended Data Fig. 5D**) to safety profile (**Figure 4D, 4E**), while leaving cytoplasmic-membrane depolarization comparatively free to vary.

Four analog leads were advanced to murine models (**Table 1**), spanning the three conditioning modes (OmegAMP-AP, OmegAMP-AT, OmegAMP-AU) and including analogs active against *A. baumannii f*or the corresponding infection model. Ω-AT-BoCo1-5 reduced *A. baumannii* ATCC 19606 MIC 32-fold from its inactive BoCo1 prototype, reaching MIC ≤ 2 μM against three multidrug-resistant Gram-negatives. Ω-AT-BoCo1-9 matched this *A. baumannii* potency gain while showing no detectable hemolysis (HC₅₀ >128 μM) and lower cytotoxicity than Ω-AT-BoCo1-5, providing a within-family safety contrast. Ω-AP-Mammutin-1-4 gained *A. baumannii* activity from the Mammutin-1 prototype (MIC 64 → 8 μM against *A. baumannii* BAA-1605) while preserving the family’s favorable hemolysis profile (HC₅₀ >128 μM). Ω-AU-GQ20-4 reached *E. coli* AIC222 at MIC = 1 μM and was the only GQ20 analog with non-detectable hemolysis (HC₅₀ >128 μM).

### Motif-guided generation combines antimicrobial activity with preserved auxiliary function

Despite antimicrobial activity being the critical aspect of AMP design, practical therapeutics could benefit from auxiliary biological functions. LPS engagement can broaden therapeutic utility by neutralizing LPS released from the outer membrane of Gram-negative bacteria, often contributing to bacterial killing through lipid A engagement^8,9^. Certain AMPs translocate across the bacterial membrane and perturb bacterial DNA, disrupting essential processes such as replication and transcription in a mechanism complementary to membrane disruption. Starting from prototypes with such auxiliary functions, we applied motif-guided generation with targeted conditioning to satisfy two objectives jointly: antimicrobial activity and preservation of auxiliary function encoded by user-specified residues mediating LPS engagement or DNA perturbation.

We selected four LPS-engaging motifs, derived from AMP prototypes, cecropin-P1^31^, LG21^32,33^, pa4^34^, and sarcotoxin-1A^35^, as well as a DNA-engaging motif, derived from bZIP^36^, a leucine zipper with no antimicrobial activity (MIC > 64 μM across all strains tested). For each motif, we generated 10 designs. The LPS-engaging designs were obtained as analogs (OmegAMP-AMT) with ≥60% sequence similarity to their prototypes, with the fixed positions corresponding to family-defining residues identified in the literature: an aromatic/hydrophobic anchor (W, GW, or GFF), clustered cationic residues, and an extended aliphatic stretch enriched in L, I, V, and A. For bZIP, family identity rests on the leucine heptad pattern rather than on discrete residues. Motif-guided generation (OmegAMP-MT) preserved this pattern, producing designs with median 30% identity to bZIP. Per family alignments are shown in **Extended Data Fig. 6A**, and full MIC, HC_50_, and CC_50_ profiles are shown in **Extended Data Fig. 6B**.

Antimicrobial screening revealed high hit rates across all five families (**Figure 5A**). All 40 LPS-engaging analogs achieved MIC ≤ 2 μM against at least one strain (39 of 40 against an MDR strain), preserving or extending the activity of their already-active prototypes. Among bZIP designs, 7 of 10 achieved MIC ≤ 2 μM and 6 reached at least one MDR strain, producing antimicrobial activity from a previously non-antimicrobial prototype. Strain-level coverage (**Figure 5B**) was once again broad against Gram-negative MDR isolates (24–88% of all designs at MIC ≤ 2 μM) and narrower against Gram-positives (≤ 20% on MRSA and vancomycin-resistant *E. faecium* ATCC 700221; 52% on *B. subtilis*).

**Figure 5.**
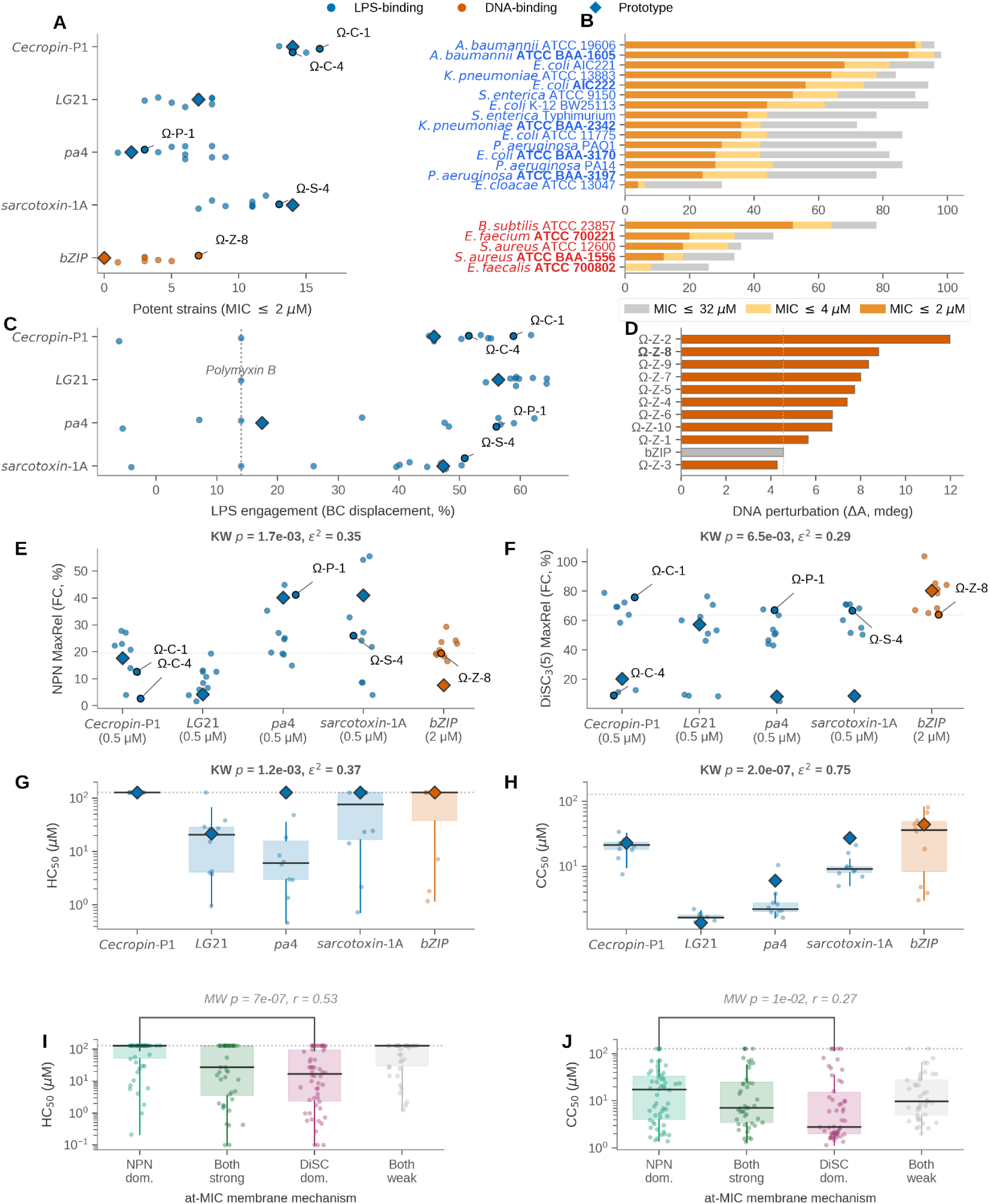
Motif-guided generation preserves biological function and adds antimicrobial activity. (A) Number of strains with MIC ≤ 2 μM per analog for the four LPS-engaging families (cecropin-P1, LG21, pa4, sarcotoxin-1A; blue) and the DNA-engaging bZIP family (orange). Diamonds, prototypes; labeled points, leads (Ω-C-1, Ω-C-4, Ω-P-1, Ω-S-4, Ω-Z-8), see also **Extended Data Fig. 6A and 6B**. (B) Fraction of motif analogs reaching MIC ≤ 32 μM (gray), ≤ 4 μM (yellow), and ≤ 2 μM (dark orange) per strain (Gram-negative, top; Gram-positive, bottom). MDR isolates in bold. (**C**) LPS engagement by BODIPY-cadaverine (BC) displacement (%) at 2 μM peptide for the four LPS-engaging families. Diamonds, prototypes; dotted vertical line, polymyxin B reference (see also **Extended Data Fig. 6C**). (**D**) DNA perturbation by CD (ΔA, mdeg) at 12.5 μM peptide for the bZIP family. Prototype in gray; lead Ω-Z-8 in bold; analogs ordered by ΔA (see also **Extended Data Fig. 6D**). (**E**, **F**) NPN (**E**) and DiSC₃(5) (**F**) MaxRel (fold-change, %) by prototype family, each measured at the per-family at-MIC concentration shown beneath the x-axis label. Diamonds, prototypes; dashed line, global median. KW p and ε² shown (see also **Extended Data Fig. 6E and 6F**). (**G**, **H**) HC₅₀ (**G**) and CC₅₀ (**H**) (μM) by prototype family. Diamonds, prototypes. KW p and ε² shown. (**I**, **J**) HC₅₀ (**I**) and CC₅₀ (**J**) (μM) grouped by at-MIC membrane-mechanism class (NPN-dominant, both-strong, DiSC-dominant, both-weak). MW p and rank-biserial r shown.

Across all four LPS-engaging families, analogs matched or exceeded their prototypes’ BC displacement, and most also exceeded polymyxin B (**Figure 5C**), showing that motif preservation did not constrain improvement. BC displacement measures peptide-LPS interaction in low-cation buffer. In serum, native divalent cations (Ca²⁺, Mg²⁺) compete with cationic peptides for LPS phosphates and attenuate engagement, which limits the clinical use of polymyxins as last-line antibiotics for carbapenem-resistant Gram-negative infections^37^. We repeated BC displacement under Ca²⁺ challenge to identify analogs whose engagement survives this regime (**Extended Data Fig. 6C**). Most analogs matched their prototype’s Ca²⁺ behavior. The pa4 prototype was Ca²⁺-suppressed, but a C-terminal E→K substitution converted 7 of 10 analogs to tolerance by removing an anionic competition site. The sarcotoxin-1A prototype was compositionally heterogeneous and its analogs spanned the full range. The prototype thus predisposes, rather than dictates, Ca²⁺ tolerance, even a Ca²⁺-suppressed prototype can yield tolerant analogs through targeted substitutions.

For bZIP, the test was whether motif-guided generation could install antimicrobial activity while preserving DNA perturbation. In the natural bZIP fold, a leucine zipper mediates dimerization while an N-terminal basic region engages DNA as an α-helix, with lysines and arginines on one face contacting the phosphate backbone^38^. The bZIP prototype used here retains the leucine-zipper pattern but carries acidic D and E residues in its N-terminal region that oppose DNA engagement. Using DNA perturbation measured by CD as a comparative readout of interaction (**Figure 5D**), nearly all analogs exceeded the prototype. The design route was visible in the alignment (**Extended Data Fig. 6A**) with analogs replacing N-terminal D and E with cationic K or R, restoring the basic region while leaving the leucine zipper intact. Position of the cationic residues mattered as well. The strongest perturbants Ω-MT-bZIP-9 and Ω-MT-bZIP-2 eliminated all anionic residues from the basic region, with cationic residues lying on the polar face of an amphipathic helix (**Extended Data Fig. 6D**). The weakest, Ω-MT-bZIP-1 and Ω-MT-bZIP-3, retained an anionic residue in the basic region on the same polar face, reducing net positive charge despite carrying higher overall cationic counts. bZIP analogs uniformly permeabilized the outer membrane (**Figure 5E**) while showing variable cytoplasmic-membrane depolarization (**Figure 5F**), consistent with one established route for intracellular-target AMPs in which outer-membrane permeabilization enables entry without full collapse of the inner-membrane potential^39^.

Membrane mechanism and safety both partitioned strongly by prototype family [NPN and DiSC₃(5), **Figure 5E**, **F**; HC₅₀ and CC₅₀, **Figure 5G, H**; all KW *p* < 0.01], reproducing the family-level pattern observed in the inactive-to-active conversion across a different and functionally distinct set of prototypes. To test whether mechanism predicts safety independently of family, we classified each peptide by its at-MIC NPN and DiSC₃(5) profile (per-family distributions in **Extended Data Fig. 6E, F**) and compared safety across the resulting classes. NPN-dominant peptides were less hemolytic (MW *p* < 0.001, *r* = –0.51; **Figure 5I**) and less cytotoxic (*p* = 0.01, *r* = –0.28; **Figure 5J**) than DiSC-dominant peptides. The two effects form a chain. Prototype family is associated with the membrane mode of action, which in turn determines mammalian-cell safety.

Five motif-guided leads illustrated the mechanism preservation enabled by this mode (**Table 1**). Among four LPS-engaging leads, Ω-AMT-cecropin-4 combined broad Gram-negative activity at MIC ≤ 2 μM, including against MDR isolates, with Ca²⁺-tolerant LPS engagement and a favorable safety window. The bZIP analog (Ω-MT-bZIP-8) acquired antibacterial activity from a non-antimicrobial leucine zipper while retaining a DNA-perturbing CD signature. We next tested whether leads from the three generation modes could reduce bacterial burden *in vivo*.

### OmegAMP leads reduce bacterial burden in topical and systemic infection models

Having established potent *in vitro* activity across all three generation modes, we asked whether this activity translates to efficacy *in vivo*, beginning with a murine skin scarification model of *A. baumannii* ATCC 19606 infection. From the wet-lab library we advanced six leads chosen to span the generation modes and to prioritize *A. baumannii* activity, HC_50_-based selectivity, broad-spectrum coverage, and activity against multidrug-resistant strains: two *de novo* designs (Ω-DP-52, Ω-DP-19), two inactive-to-active analogs of the BoCo1 template (Ω-AT-BoCo1-5, Ω-AT-BoCo1-9), and two motif-guided analogs preserving the cecropin-P1 LPS-engaging motif (Ω-AMT-cecropin-1, Ω-AMT-cecropin-4). Parent templates were not carried into the animal experiments, as for the inactive-to-active family the *in vitro* MIC contrast already establishes the activity gain, and for the active motif templates the objective was to validate the analogs themselves rather than to demonstrate strict improvement, which also spared animals. A superficial dorsal wound was abraded and inoculated, peptides were applied topically one hour after infection, and bacterial burden in the excised wound was quantified two and four days post-infection, with polymyxin B as a benchmark antibiotic control (**Figure 6A**).

**Figure 6.**
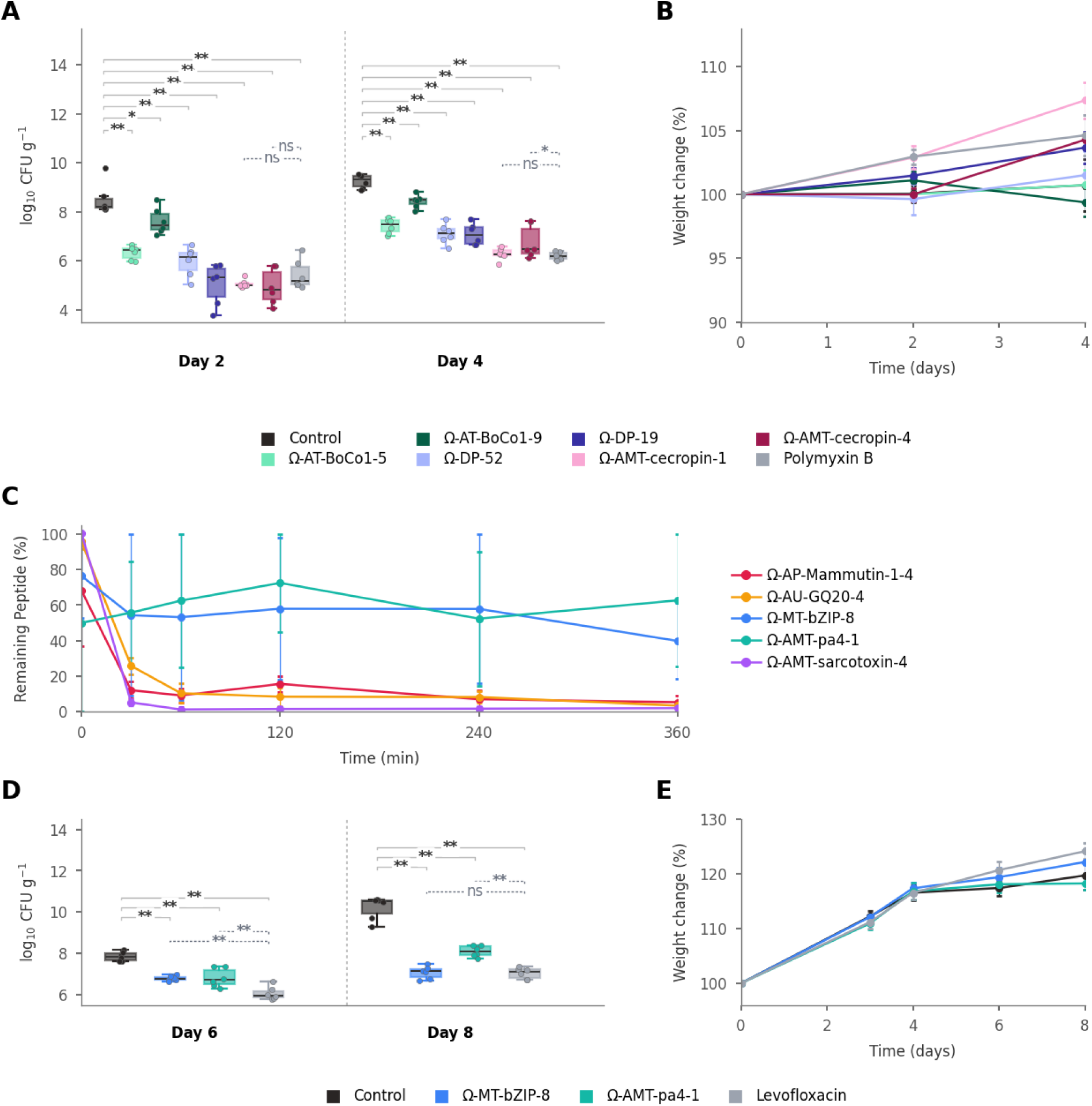
OmegAMP-designed leads reduce *Acinetobacter baumannii* bacterial burden in murine infection models. **(A)** Bacterial burden (log₁₀ CFU g⁻¹) in a murine skin scarification model of *A. baumannii* ATCC 19606 infection, at days 2 and 4 post-infection. Six-week-old female CD-1 mice were treated topically at 10×MIC 1h after infection. Groups span generation modes: inactive-to-active BoCo1 analogs (Ω-AT-BoCo1-5, Ω-AT-BoCo1-9), de novo leads (Ω-DP-52, Ω-DP-19), and motif-guided cecropin-P1 analogs (Ω-AMT-cecropin-1, Ω-AMT-cecropin-4), with untreated control and polymyxin B as references. Boxes, interquartile range and median; whiskers, range; points, individual mice (*n = 6* per group per timepoint). Solid brackets: significance versus control by two-sided exact Mann-Whitney tests, with a Kruskal-Wallis test across all groups in each day-block. Dashed gray brackets: the motif-guided analogs versus the reference antibiotic (polymyxin B). Significance levels: * *p* < 0.05, ** *p* < 0.01, *** *p* < 0.001; ns, not significant. **(B)** Body-weight change (%, relative to day 0) over the skin scarification study for the groups in (A). Points, mean; error bars, SEM. **(C)** Proteolytic stability of five leads incubated with protease, reported as remaining intact peptide (%) over 6 h (Ω-AP-Mammutin-1-4, Ω-AU-GQ20-4, Ω-MT-bZIP-8, Ω-AMT-pa4-1, Ω-AMT-sarcotoxin-4). Points, mean of two replicates; error bars, range. **(D)** Bacterial burden (log₁₀ CFU g⁻¹) in a neutropenic murine thigh infection model of A. baumannii ATCC 19606, at days 6 and 8 post-infection. Female CD-1 mice were rendered neutropenic with cyclophosphamide, infected by intramuscular injection into the thigh, and treated intraperitoneally at 10×MIC 2 h after infection. Groups: Ω-MT-bZIP-8 and Ω-AMT-pa4-1, with untreated control and levofloxacin as references. Conventions and statistics as in (A). **(E)** Body-weight change (%) over the thigh infection study for the groups in (D). Conventions as in (B).

All six designs reduced wound burden relative to the untreated control, and the two motif-guided cecropin-P1 analogs were the most efficacious at both time points (**Figure 6A**). At day 2, Ω-AMT-cecropin-4 and Ω-AMT-cecropin-1 reduced burden by 3.4 and 3.2 log₁₀, respectively (exact Mann-Whitney, *p < 0.01* versus the untreated control), comparable to polymyxin B (3.0 log₁₀), a last-resort therapy for *A. baumannii* infections, from which neither differed significantly. The *de novo* lead Ω-DP-19 was similarly effective (2.9 log₁₀, *p < 0.01*). The remaining designs also reduced burden significantly, by 2.0 log₁₀ (Ω-DP-52), and 1.8 log₁₀ (Ω-AT-BoCo1-5), both at *p < 0.01*, and by 0.8 log₁₀ (Ω-AT-BoCo1-9) at *p < 0.05*, although the two BoCo1 analogs resulted in significantly higher burden than polymyxin B (*p < 0.05*). This ordering was preserved at day 4, where every design was again significant against the control burden (*p < 0.01*). Ω-AMT-cecropin-1 remained comparable to polymyxin B (3.0 versus 3.2 log₁₀, not significant), whereas Ω-AMT-cecropin-4 (2.9 log₁₀) produced a significantly smaller reduction than polymyxin B (*p < 0.05*). The peptides were well tolerated. No treatment group lost body weight over the four-day course, with individual animals remaining between 91% and 114% of their starting weight and group means holding at or slightly above baseline (**Figure 6B**).

We next asked whether the leads would remain active in a systemic model, where a peptide is exposed to serum proteases rather than applied topically. Because proteolytic stability is a principal obstacle to peptide therapeutics and a prerequisite for systemic exposure, we used it as a selection filter ahead of the more demanding thigh infection model. A panel of five designed peptides spanning the analog and motif-guided modes was incubated with protease and the fraction of intact peptide was tracked over six hours (**Figure 6C**). Most candidates degraded rapidly, falling below 10% intact within one hour, whereas Ω-AMT-pa4-1 and Ω-MT-bZIP-8 were markedly more stable, retaining approximately 63% and 40% of intact peptide at six hours. These two peptides, representing the LPS-engaging and DNA-perturbing motif-guided modes, were advanced to the neutropenic thigh infection model.

In the neutropenic thigh model, *A. baumannii* ATCC 19606 was established in the muscle of cyclophosphamide-immunosuppressed mice, peptides were administered systemically, and bacterial burden was quantified at days six and eight, with levofloxacin as the antibiotic control (**Figure 6D**). Each peptide was compared both against the untreated control, to test efficacy, and against levofloxacin, to test whether it matched standard-of-care. By day 8, both designs had significantly reduced the thigh bacterial load relative to the untreated control, Ω-MT-bZIP-8 by 3.4 log₁₀ and Ω-AMT-pa4-1 by 2.4 log₁₀ (*p < 0.01*). Ω-MT-bZIP-8 achieved the same 3.4 log₁₀ reduction as levofloxacin and did not differ significantly from it, indicating comparable activity to a fluoroquinolone antibiotic in this systemic model, whereas Ω-AMT-pa4-1 was significantly less effective than levofloxacin (*p < 0.01*). At the earlier day-6 timepoint both peptides had already reduced burden significantly relative to control, by approximately 1.1 log₁₀ each (*p < 0.01*), although both were significantly less effective than levofloxacin at that point (*p < 0.01*). Both peptides were well tolerated, with no weight loss over the eight-day course (**Figure 6E**). The systemic efficacy of Ω-MT-bZIP-8, a proteolytically stable analog that acquired antimicrobial activity from a non-antimicrobial leucine-zipper prototype while retaining DNA perturbation, demonstrates that motif-guided OmegAMP designs can combine multi-mechanism function with the stability required for use beyond a topical setting.

## Discussion

OmegAMP delivers conditional AMP generation with explicit control over net charge, mean hydrophobicity, and length, together with preservation of user-defined sequence motifs that encode auxiliary functions. A single conditional diffusion model integrates these capabilities and supports *de novo*, analog, and motif-guided generation. Wet-lab evaluation across all three generation modes returned high hit rates of potent antimicrobials, including against MDR Gram-negative isolates.

As hypothesized, analogs followed prototype properties beyond those we directly conditioned on. Across the analog set, the prototype family accounted for substantial variance in outer-membrane permeabilization, hemolysis, and cytotoxicity. This inheritance follows directly from the analog design. Because analogs retain the sequence identity of their prototype, composition, length, and cationic-amphipathic architecture are conserved within a family, so amphipathic arrangement of residues around the helix and the placement of residues driving membrane partitioning remain largely fixed even as net charge and mean hydrophobicity shift. The mode of membrane disruption then sets mammalian-cell safety. Peptides with outer-membrane-dominant disruption at MIC against *A. baumannii* were substantially less hemolytic and less cytotoxic than those with cytoplasmic-dominant disruption. Mammalian cells lack the anionic outer membrane of Gram-negative bacteria but share their cytoplasmic bilayer. A peptide with outer-membrane-dominant disruption therefore has no corresponding target in a mammalian cell, and because outer-membrane permeabilization alone can be sufficient for antibacterial activity, this selectivity costs no potency^9,40^. Gram-positive bacteria, lacking an outer membrane, are susceptible only through the cytoplasmic membrane, and accordingly Gram-positive activity is tracked with hemolysis and cytotoxicity. Screening candidate prototypes for outer-membrane-dominant disruption before analog generation is therefore a tractable route to wider safety windows, though it also limits Gram-positive coverage.

The same membrane interfaces also explain the strain-level susceptibility pattern across our bacterial panel, since each of the major resistance mechanisms targets the layer where the corresponding AMPs act. Among the Gram-negatives, *A. baumannii* BAA-1605 was the most susceptible MDR isolate, consistent with the clinical use of polymyxins, cationic peptides themselves, as last-resort therapy for *A. baumannii*. Carbapenem resistance arises from carbapenemase activity, a route distinct from the lipid A modification or LPS loss required for polymyxin resistance^41^, so the strain has no pre-existing defense against peptides targeting the LPS layer. *K. pneumoniae* BAA-2342 was the most resistant. Its outer polysaccharide capsule shields the outer membrane from cationic peptides, a defense independent of its carbapenemase^42^, and places the first barrier well before the LPS layer that NPN-dominant peptides engage. Among Gram-positives, the gap between *B. subtilis* and the four pathogenic isolates is consistent with cytoplasmic-membrane defenses that pathogens deploy against cationic peptides. Positively charged groups added to membrane lipids (*mprF*) and teichoic acids (*dltABCD*), reduce the net negative surface charge, and secreted peptidases degrade the peptides directly^43^. These defenses help explain the limited Gram-positive coverage across our generation modes. The mechanism-safety analysis above accounts for why the peptides that do overcome them tend toward higher mammalian toxicity.

Motif-guided generation tested whether a residue-level functional constraint and an antimicrobial objective could be satisfied jointly. Across all four LPS-engaging families, analogs reached potent antimicrobial activity while preserving or improving LPS engagement relative to their prototypes, and the cecropin-derived lead retained engagement under Ca²⁺ challenge. The bZIP case is more striking. Redesigned non-antimicrobial leucine zipper acquired antimicrobial activity while retaining DNA perturbation. Motif-guided generation can therefore add antimicrobial activity onto a peptide whose primary function is non-antimicrobial, while preserving that primary function. These motif-guided leads were also the most efficacious *in vivo*. The cecropin-derived analogs reduced *A. baumannii* burden in a murine skin infection to the level reached by polymyxin B, and Ω-MT-bZIP-8, which resisted proteolysis, matched the systemic efficacy of levofloxacin in a neutropenic thigh infection. Motif-guided generation therefore yields peptides that combine multi-mechanism function with therapeutically relevant potency, stability, and safety.

Several limitations define the scope of these conclusions. Analog generation enforced a ≥60% sequence-similarity floor, so the family-level associations are established only above it. The bZIP designs, at roughly 30% identity to their prototype, hint that these associations persist at lower similarity, though we did not test this systematically. The membrane-mechanism readouts are narrower still. NPN and DiSC₃(5) were recorded at MIC against a single strain, *A. baumannii* ATCC 19606, leaving open whether the mechanism-safety relationship extends above MIC or to Gram-positive organisms, which lack the LPS outer membrane the NPN assay reports on. Those organisms were also poorly covered across all three generation modes. The model itself is deliberately partial. It conditions on net charge, length, and mean hydrophobicity, but not on local features such as residue spacing and hydrophobic-face geometry, which our secondary-structure data show also shape potency, and it optimizes for activity without conditioning on toxicity. The 20-strain panel omits slow-growing and intracellular pathogens, and the *in vivo* work covers only skin and thigh infection.

OmegAMP can be extended for further controllability without architectural changes. The conditioning vector can accommodate additional design objectives alongside property targets, with toxicity, proteolytic stability, and synthesizability as immediate candidates. Resistance propensity is a further candidate of particular interest, since OmegAMP’s existing controls over charge and hydrophobicity directly engage physicochemical features empirically associated with reduced resistance evolution in AMPs^6^. The same framework extends in principle to other classes of bioactive peptides whose function is governed by sequence-derived physicochemical properties, and to design problems requiring simultaneous satisfaction of distinct objectives. Controllable generation may therefore provide a general route from sequence sampling toward hypothesis-driven molecular design.

## Corresponding author

Requests for further information and resources should be directed to and will be fulfilled by the lead contact, Ewa Szczurek.

## Data availability

Wet-lab characterization data together with the designed peptide sequences and accompanying metadata have been deposited at Zenodo and are publicly available as of the date of publication. The DOI is listed in the key resources table. This paper also analyzes existing, publicly available data from DBAASP, dbAMP, DRAMP, UniProtKB, APD, and Peptipedia; accession URLs are listed in the key resources table.

## Code availability

All original code, including the OmegAMP framework and trained model weights, has been deposited at Zenodo (https://doi.org/10.5281/zenodo.21821343) and is publicly available as of the date of publication. The generative framework and inference code are mirrored at https://github.com/szczurek-lab/OmegAMP, and the analysis and figure-reproduction code at https://github.com/szczurek-lab/omegamp-dashboard.

The interactive dashboard is available at https://szczurek-lab.github.io/omegamp-dashboard. Any additional information required to reanalyze the data reported in this paper is available from the lead contact upon request.

## Acknowledgments

This project was supported by the European Research Council (ERC) under the European Union’s Funding Horizon 2020 research and innovation programme (grant agreement No. 101125506 – DOG-AMP to E.S). C.F.-N. holds a Presidential Professorship at the University of Pennsylvania and acknowledges funding from the National Institute of General Medical Sciences of the National Institutes of Health under award number R35GM138201 and the Defense Threat Reduction Agency (DTRA; HDTRA1-21-1-0014). S.J. and D.S. acknowledge funding from the Alexander von Humboldt foundation.

The Graphical Abstract and Figure 1 were created at https://BioRender.com.

## Author Contributions

**Conceptualization,** P.S., M.D.T.T., D.S., L.H., C.d.l.F.-N., and E.S.; **Methodology,** P.S., M.D.T.T., D.S., L.H., B.P.-S and E.S.; **Software,** P.S., D.S., and L.H.; **Formal Analysis,** P.S.; **Investigation,** P.S. (computational) and M.D.T.T. (experimental); **Validation,** M.D.T.T.; **Data Curation,** P.S. and M.D.T.T.; **Visualization,** P.S.; **Writing – Original Draft,** P.S., M.D.T.T., D.S., L.H., C.d.l.F.-N., and E.S.; **Writing – Review & Editing,** all authors; **Supervision,** S.J, S.G., F.J.T., C.d.l.F.-N., and E.S.; **Project Administration,** C.d.l.F.-N. and E.S.; **Funding Acquisition,** S.J, S.G., F.J.T., C.d.l.F.-N., and E.S.; **Resources,** C.d.l.F.-N. and E.S.

## Declaration of Generative AI and AI-Assisted Technologies

During the preparation of this work, the authors used Claude (Anthropic) to assist with language editing and consistency checking of the manuscript, to build the interactive dashboard, and to support data analysis and figure generation, and ChatGPT (OpenAI) to assist with language editing and manuscript organization. After using these tools, the authors reviewed and edited the content as needed and take full responsibility for the content of the publication.

## Supplementary Information

**Data S1.** *In silico* prototype set, related to Extended Data Fig. 2.

Excel workbook with one sheet each for the 500 active and 500 inactive prototype peptide sequences sampled from DBAASP and used to benchmark OmegAMP analog generation against HydrAMP. Each sheet lists the DBAASP identifier and the peptide sequence.

**Data S2.** OmegAMP wet-lab characterization dataset, related to Figures 3, 4, 5, and 6.

Excel workbook containing one sheet per measurement type for the 215 wet-lab characterized peptides: (i) peptide metadata (sequence, generation mode, conditioning strategy, prototype, design objective and round); (ii) MIC matrix across the 20-strain panel; (iii) HEK293T CC₅₀; (iv) human-RBC HC₅₀; (v) NPN MaxRel and AUC, measured both at each peptide’s MIC against *A. baumannii* ATCC 19606 and at a family-anchored fixed concentration; (vi) DiSC₃(5) MaxRel and AUC under the same two regimes; (vii) BODIPY TR cadaverine displacement in triplicate, at 2, 8 and 32 μmol L⁻¹ without added Ca²⁺ and at 32 μmol L⁻¹ under Ca²⁺ charge-shielding; (viii) DNA CD perturbation of the bZIP series (ΔA at 12.5, 25 and 50 μmol L⁻¹ peptide and ΔA_max in mdeg; positive– and negative-band shifts Δλ in nm); (ix) BeStSel secondary-structure deconvolutions across the four CD solvent conditions (H₂O, TFE/H₂O 3:2 v/v, 10 mM SDS/H₂O, MeOH/H₂O 1:1 v/v); (x) proteolytic stability (percentage of intact peptide over 6 h of protease exposure, two replicates); (xi) *in vivo* efficacy in the murine skin-scarification and neutropenic-thigh A. baumannii ATCC 19606 models, as per-mouse bacterial burden (CFU g⁻¹, days 2 and 4 or 6 and 8 respectively) and body weight relative to day 0; and (xii) maximum sequence identity of each de novo peptide to the generative training set. Antibiotic controls (polymyxin B, levofloxacin), the Triton X-100 lysis control and the untreated infection control are included wherever they were measured alongside. MIC values reported as >64 μmol L⁻¹ and CC₅₀/HC₅₀ values reported as >128 μmol L⁻¹ are right-censored at the highest concentration tested. A README sheet documents every sheet, column and unit.

## Methods

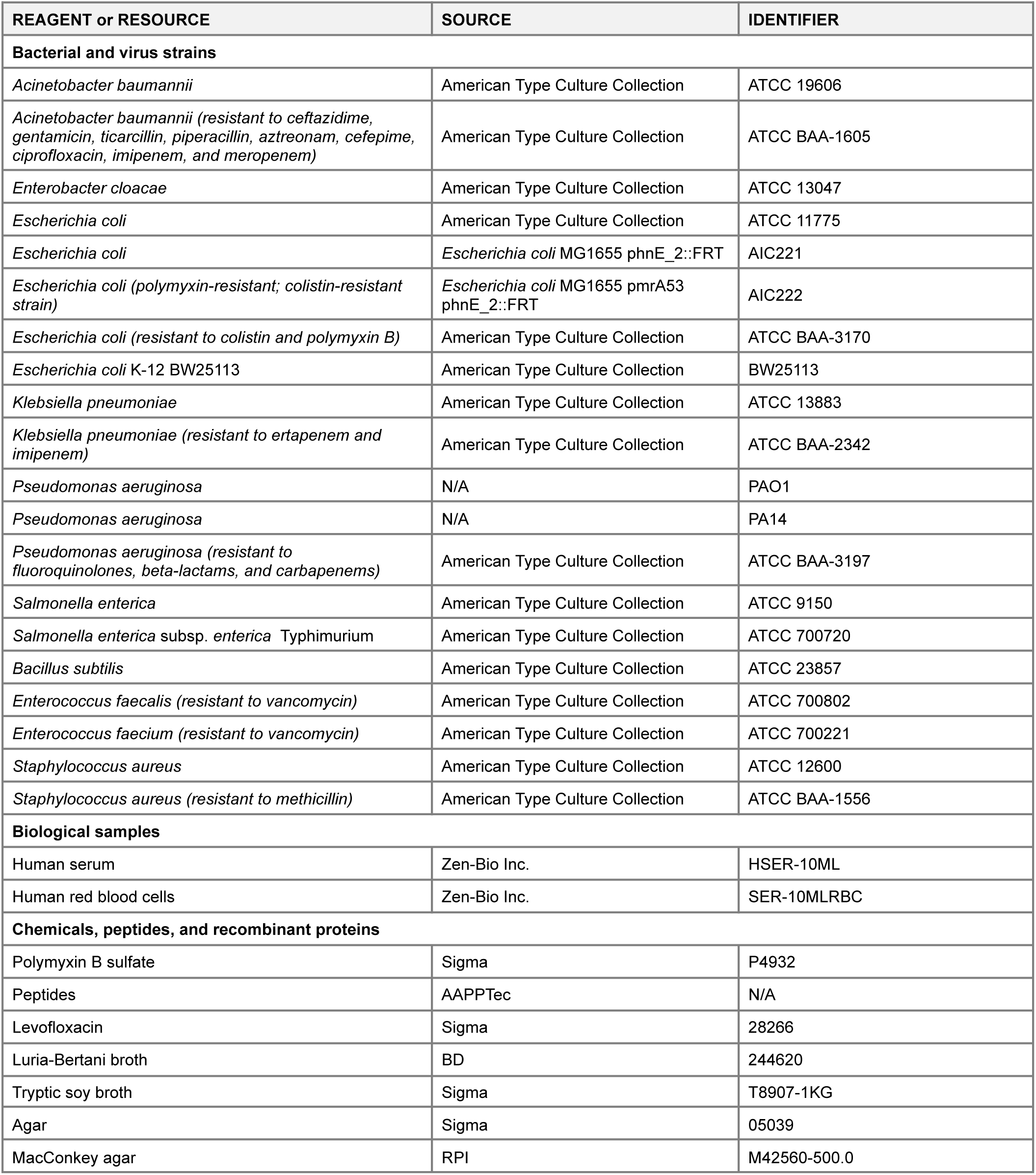
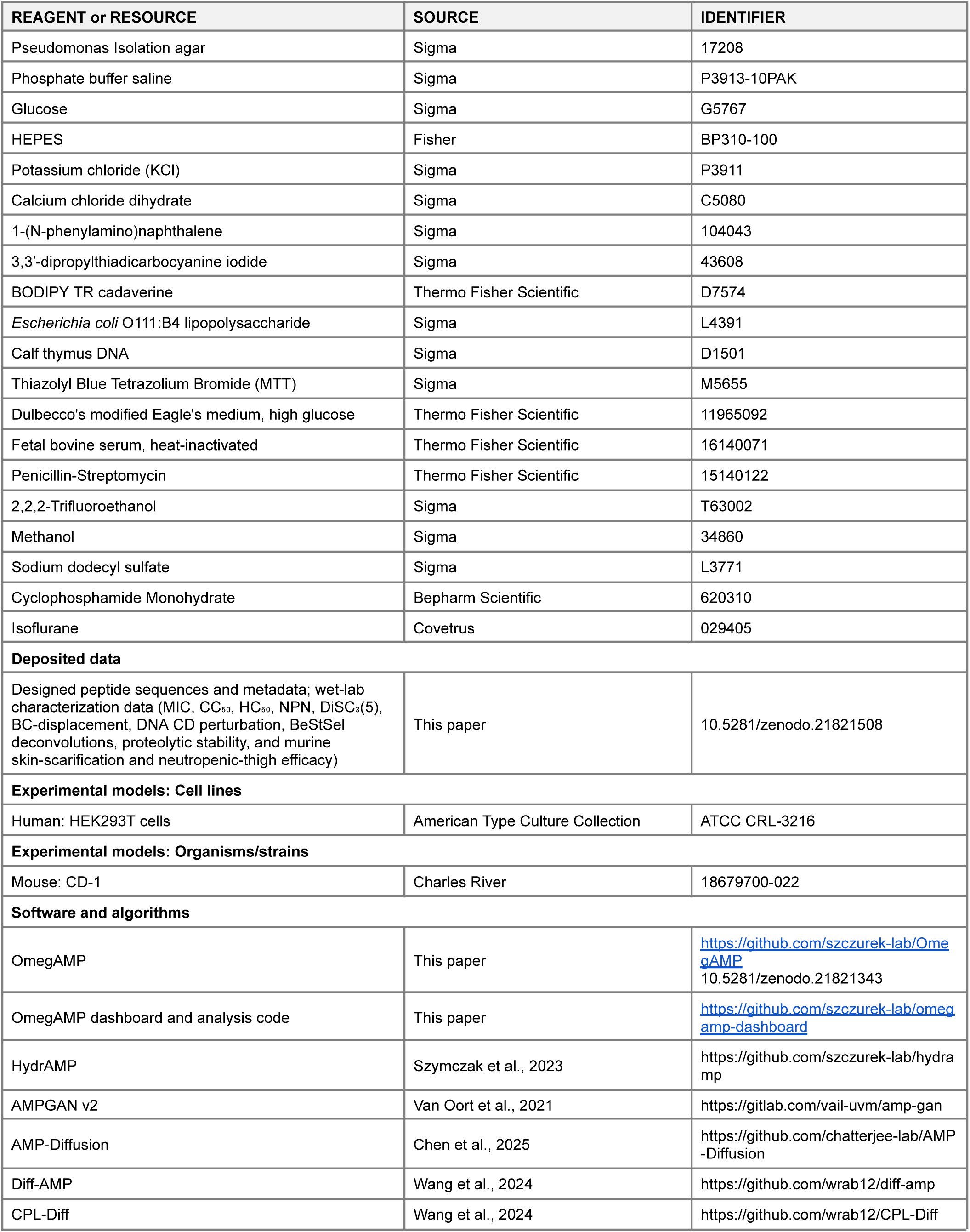
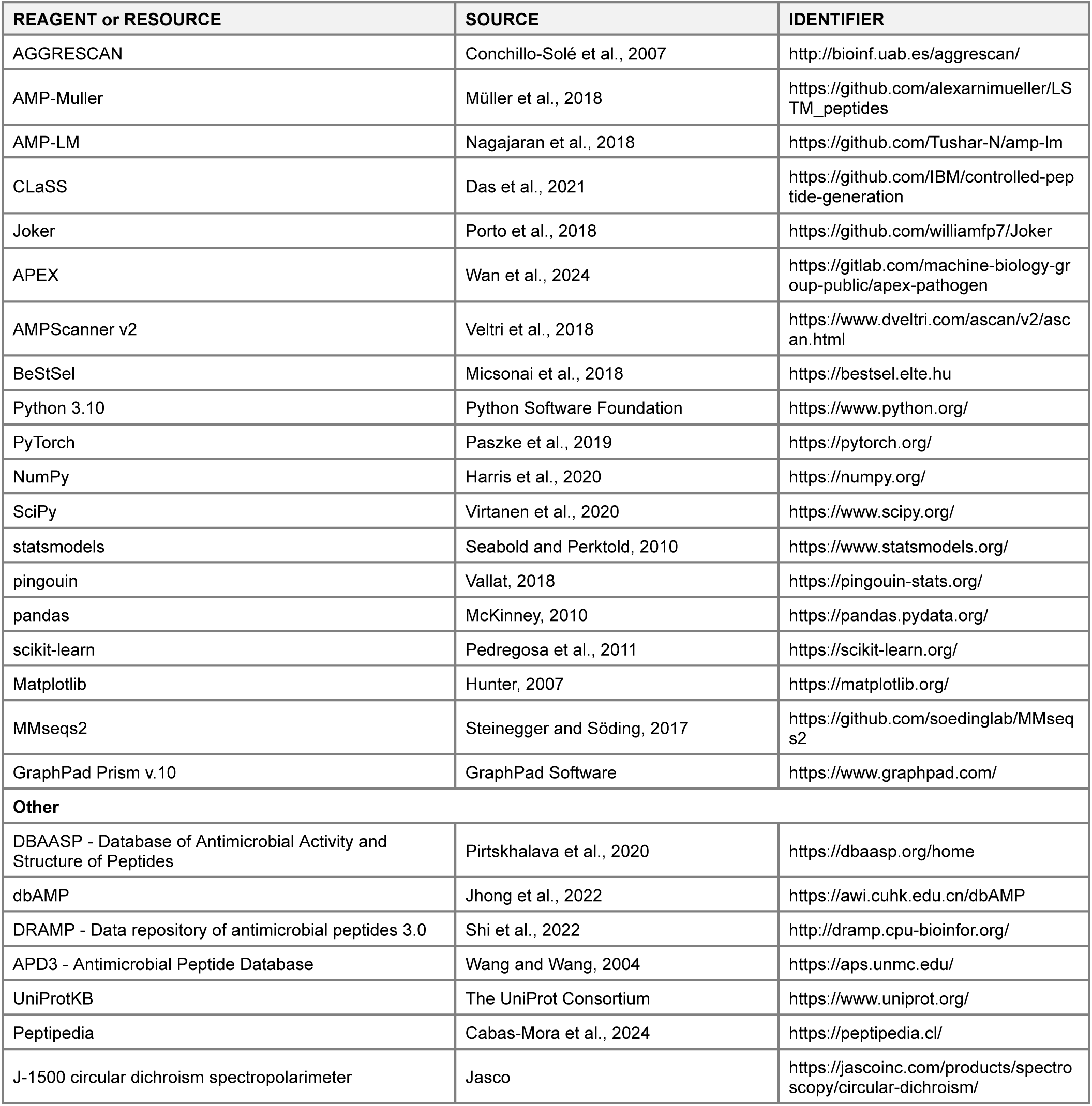
Key Resources Table.

## Experimental Model and Study Participant Details

### Bacterial strains and growth conditions

Bacterial strains used in this study included *Acinetobacter baumannii* ATCC 19606, *A. baumannii* ATCC BAA-1605 (carbapenem-resistant), *Enterobacter cloacae* ATCC 13047, *Escherichia coli* ATCC 11775, *E. coli* AIC221 [MG1655 phnE_2::FRT; polymyxin-sensitive control], *E. coli* AIC222 [MG1655 pmrA53 phnE_2::FRT; polymyxin-resistant], *E. coli* ATCC BAA-3170 (carbapenem-resistant), *E. coli* K-12 BW25113, *Klebsiella pneumoniae* ATCC 13883, *K. pneumoniae* ATCC BAA-2342 (carbapenem-resistant), *Pseudomonas aeruginosa* PAO1, *P. aeruginosa* PA14, *P. aeruginosa* ATCC BAA-3197 (fluoroquinolone-resistant), *Salmonella enterica* ATCC 9150, *S. enterica* subsp. *enterica* Typhimurium ATCC 700720, *Bacillus subtilis* ATCC 23857, *Staphylococcus aureus* ATCC 12600, *S. aureus* ATCC BAA-1556 (methicillin-resistant), *Enterococcus faecalis* ATCC 700802 (vancomycin-resistant), and *Enterococcus faecium* ATCC 700221 (vancomycin-resistant). *Pseudomonas* isolates were cultured on Pseudomonas Isolation Agar, whereas all other bacterial strains were propagated in Luria-Bertani (LB) agar or broth. For each experiment, cultures were initiated from a single colony, grown overnight at 37 °C, and diluted 1:100 into fresh medium before growth to mid-logarithmic phase.

### Eukaryotic cells culture

Human embryonic kidney (HEK293T) cells were obtained from the American Type Culture Collection (CRL-3216). Cytotoxicity toward mammalian cells was assessed using HEK293T cells and the MTT viability assay. The cells were cultured in high-glucose Dulbecco’s modified Eagle’s medium supplemented with 1% penicillin and streptomycin (antibiotics) and 10% fetal bovine serum and grown at 37 °C in a humidified atmosphere containing 5% CO_2_.

### Skin scarification infection mouse model

Six-week-old female CD-1 mice were anesthetized, and the dorsal skin was shaved and superficially abraded with a needle to generate a linear wound. *A. baumannii* ATCC 19606 was grown in LB medium to an OD₆₀₀ of 0.5, washed twice with sterile PBS (pH 7.4) by centrifugation at 9,391 × g for 2 min, and resuspended to 6.66 × 10⁵ CFU mL⁻¹. A 20 μL aliquot of the bacterial suspension was applied directly to the scarified area. Peptides were diluted in sterile water and administered topically to the infected wound at 10×MIC values 1 h post-infection. Two and four days post-infection, mice were euthanized, and a uniform section of scarified skin was excised, homogenized using a bead beater at 25 Hz for 20 min, 10-fold serially diluted, and plated on MacConkey agar for CFU quantification. Experiments were performed with six mice per group. Mice were single-housed to minimize cross-contamination and maintained under a 12 h light/dark cycle at 22 °C and 50% humidity. All animal procedures were reviewed and approved by the University Laboratory Animal Resources (ULAR) at the University of Pennsylvania under protocol 806763.

### Thigh infection mouse model

Six-week-old female CD-1 mice were rendered neutropenic by intraperitoneal administration of cyclophosphamide at 150 mg kg⁻¹ and 100 mg kg⁻¹ on days 0 and 3 before infection (day 4), respectively. *A. baumannii* ATCC 19606 was grown in LB broth, washed twice with sterile PBS, and diluted in PBS to 9.66 × 10⁵ CFU mL⁻¹. Mice were infected by intramuscular injection of 100 μL of the bacterial suspension into the right thigh. Peptides were administered intraperitoneally 2 h post-infection at 10×MIC. Two days post-infection, mice were euthanized, and a uniform section of tissue from the infected right thigh was excised, homogenized using a bead beater at 25 Hz for 20 min, 10-fold serially diluted, and plated on MacConkey agar for CFU quantification. Experiments were performed with six mice per group. Mice were housed in groups of three and maintained under a 12 h light/dark cycle at 22 °C and 50% humidity. All animal procedures were reviewed and approved by the University Laboratory Animal Resources (ULAR) at the University of Pennsylvania under protocol 807055.

## Method Details

The procedures below describe the OmegAMP generative framework, the *in silico* experiments, the design of the peptide library advanced to experimental characterization, and the wet lab procedures. A full mathematical formulation of the diffusion framework and of the training and inference algorithms is provided in the OmegAMP paper^44^. MIC cutoffs applied to DBAASP records, the ≤ 32 and ≥ 128 values that define active and inactive peptides, are in μg/mL as reported in DBAASP; all other MIC values in this work are in μM.

### Training dataset

The generative model was trained on 810,283 peptides of length ≤ 100 residues, assembled from two sources. The curated antimicrobial peptide subset (35,878 sequences) was aggregated from the AMPScanner v2 training set^45^, dbAMP 2.0^46^ and DRAMP 3.0^47^ with exact duplicates removed across sources; these curated sequences carry a positive antimicrobial-activity label. The general peptide subset (774,405 sequences) was sampled from Peptipedia^48^, restricted to functional peptides extracted from UniProtKB^49^, that Peptipedia’s prediction pipeline classified as Antibacterial, Anti-Gram-positive or Anti-Gram-negative; these sequences carry a negative antimicrobial-activity label as these sequences are not guaranteed to be antimicrobial. The general peptide subset was included to encourage the model to better learn the distribution of functional and chemically stable peptides close to antimicrobials. Furthermore, it offers additional examples of the mapping between charge, hydrophobicity, and length requirements, thereby improving the controllable capabilities of the model.

### Peptide embedding

Each peptide is represented in a continuous 6-dimensional embedding space rather than directly on discrete amino-acid sequences. For a peptide sequence *s*, each residue is mapped to a 6-dimensional vector through five physicochemically-derived amino-acid scales and a binary presence flag (1 for a residue, 0 for a padding position): hydrophobicity on the Wimley-White interfacial scale^50^, the isoelectric point (pI) of each amino acid^51^, α-helix propensity on the Levitt scale^52^, transmembrane tendency on the Zhao-London scale^53^ and the average amino-acid selectivity index AASI^54^. The five scales are jointly injective, so the map from amino acid to its 6-dimensional embedding vector is reversible. When the reverse trajectory terminates, each position of the denoised embedding is decoded to the amino acid (or padding token) whose 6-dimensional reference vector is closest in Euclidean (L2) distance, and decoding stops at the first padding token.

### Property profile vector

Conditioning information is supplied as a 4-dimensional property profile vector *c* = (*A*, *C*, *L*, *H*), where *A* ∈ {0, 1} is the antimicrobial activity indicator (1 for sequences from curated AMP databases, 0 otherwise), *C* is the net charge at physiological pH, *L* is the sequence length and *H* is the mean hydrophobicity on the Eisenberg scale. *C*, *L* and *H* are computed deterministically from the sequence, avoiding dependence on classifier-derived labels.

### Denoising network architecture

The denoising network is a one-dimensional U-Net^55^ with skip connections and additional attention layers, operating on the 6 × *100* (100 being the maximum sequence length) embedding tensor *E*. The conditioning vector *c* is encoded by a dense layer and injected by concatenation at multiple network stages so that target properties influence denoising throughout the trajectory. Full architectural details are provided in the OmegAMP paper^44^.

### Conditional training

OmegAMP was trained as a denoising diffusion probabilistic model^56^ with a cosine variance schedule^57^. The training objective was the mean squared error between the predicted and ground-truth velocity (the v-prediction parameterization^58^), computed under stochastic masking of the conditioning vector. The activity indicator A was always retained. For each training example, the number of masked entries among C, L, and H was drawn uniformly from {0, 1, 2, 3}, and that many entries were then masked at random. As a result, the model is trained on fully specified, partially specified, and fully masked profiles, so that at inference any subset of C, L, and H can be fixed while the remaining entries stay masked. Hyperparameters (optimizer, learning rate, batch size, number of training steps) are reported in the OmegAMP paper^44^.

### Conditioning strategies

At inference, *c* is populated under one of three strategies. **OmegAMP-U** (unconditional): all entries except *A* are masked. **OmegAMP-T** (targeted): point values or intervals are supplied for any subset of *C*, *L*, *H*; explicit values populate *c* directly, and for an interval (e.g., *C* ∈ [2, 10]) a target value is drawn uniformly from the interval once per generated peptide. **OmegAMP-P** (prototype-derived): *c* is computed from a prototype peptide, with the relaxation parameter σ ∈ [0, 1] setting how far each entry may deviate from the prototype value. At σ = 0, *C*, *L* and *H* match the prototype exactly; at σ = 1, each may deviate by up to one training-set standard deviation. *A* is never masked or relaxed.

### Generation modes

The reverse diffusion trajectory is initialized under one of three modes. **OmegAMP-D** (*de novo*): the trajectory starts from pure Gaussian noise at *t* = *T*. **OmegAMP-A** (analog): the prototype is encoded into *E*, forward-diffused to *t* = τ · *T* (τ ∈ [0, 1]), and reverse diffusion proceeds from this partially noised embedding; τ = 0 reproduces the prototype, τ → 1 approaches the *de novo* regime. **OmegAMP-M** (motif-guided): a template fixes amino acid residues at designated positions. At each reverse step, reconstruction guidance takes a gradient-descent step on the squared Euclidean distance between the denoised embedding and the target residues’ reference embeddings at the fixed positions, while the non-fixed positions denoise freely. Modes and conditioning strategies combine modularly, giving OmegAMP-DU, –DT, –DP; –MU, –MT, –MP; –AU, –AT, –AP; and –AMU, –AMT, –AMP.

### APEX antimicrobial activity predictor

APEX^22^ is a neural-network ensemble of 34 strain-specific MIC predictors. For each generated peptide, the per-strain predicted MIC values (in μM) were aggregated to a single per-peptide score by taking the minimum across the 34 strains. When summarizing sets of peptides (e.g., per grid coordinate in the conditioning sweep, per prototype in the analog sweep), MIC_APEX_ was further aggregated across peptides in the set by arithmetic mean of log_2_ MIC_APEX_ or by median, as indicated in the corresponding figure legend.

### Aggregation propensity

Per-sequence aggregation propensity was computed with AGGRESCAN^59^. Each residue was assigned its AGGRESCAN aggregation-propensity value, the values were averaged within a sliding window of five residues, and the mean of these window averages was the per-peptide score (for peptides shorter than five residues, the mean over all residues was used).

### Property landscape of characterized AMPs

The experimental MIC and aggregation-propensity landscapes were computed from n = 5,207 experimentally characterized peptides aggregated from DBAASP. For each peptide, MIC₅₀ is the median MIC across the evaluated strain set, corresponding to the 20-strain panel in the wet-lab experiments and to the strains reported in DBAASP in the computational evaluation. Aggregation propensity was scored with Aggrescan. Spearman rank correlation coefficients between each property (charge, length, hydrophobicity) and MIC₅₀ were computed, together with partial Spearman coefficients controlling for the remaining two properties. To assess how well these properties predict potency, five-fold cross-validated ridge linear regression and random forest models were trained to predict log₂ MIC₅₀ from physicochemical descriptors (charge, hydrophobicity, length), amino acid composition, or both, and evaluated by Spearman ρ and the coefficient of determination (R²).

### Conditioning accuracy

Conditioning accuracy was assessed by sampling under targeted (T) conditioning and computing the fraction of generated peptides whose realized properties fall within ± 0.25 units of the target charge, ± 0.1 units of the target mean hydrophobicity and the exact target length. For charge and hydrophobicity these tolerances are stricter than the change induced by a single residue substitution; length was required to match the target exactly. For each property pair (*C* × *H*, *C* × *L*, *H* × *L*), a two-property deviation map was generated over 49 target coordinates on a regular grid (*C* ∈ {0, 2, 4, 6, 8, 10, 12}, *H* ∈ {-0.75, –0.5, –0.25, 0, 0.25, 0.5, 0.75}, *L* ∈ {5, 10, 15, 20, 25, 30, 35}), with 500 peptides per coordinate. Separately, the root-mean-square error (RMSE) between target and realized values was computed per property under single-property and three-property conditioning.

### Conditioning benchmark vs. baselines

OmegAMP-T, OmegAMP-P and OmegAMP-U were compared to seven published AMP generative models: AMP-GAN^12^, HydrAMP^11^, Diff-AMP^60^, AMP-Diffusion^18^, AMP-Muller^61^, AMP-LM^62^ and CPL-Diff^63^. OmegAMP, HydrAMP and CPL-Diff samples (50,000 sequences per method) were generated locally with each method’s publicly released code. For the remaining baselines, pre-computed samples deposited with the original publications were used. The fraction of generated peptides whose realized *C*, *L* and *H* fell within the therapeutically relevant range *C* ∈ [2, 10], *L* ∈ [5, 30], *H* ∈ [-0.5, 0.8] was reported per method under single-property, pairwise and three-property regimes; for baselines lacking explicit property control, this fraction scores their default sampling distribution against the target range. Experimental success rate was computed as the cumulative fraction of tested peptides reaching MIC at or below a given threshold against at least one strain, and compared with previously reported peptides from AMP-Diffusion, HydrAMP, CLaSS and Joker^11,17,18,26^, using the MIC values reported in those studies.

### Sequence similarity

Pairwise sequence similarity between two peptides was quantified as a normalized local-alignment bitscore computed with MMseqs2^64^ in easy-search mode. For a query *s* aligned against a target *t*, the similarity score was defined as

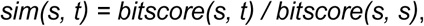

the local-alignment bitscore of *s* against *t* divided by the self-alignment bitscore of *s* against itself. The score lies in [0, 1] and is 1 when *t* contains *s* as an exact substring. Alignments used an AMP-derived substitution matrix derived in this work by aligning sequences in the curated AMP training set; the matrix file is deposited with the code repository. MMseqs2 was run with non-default settings reflecting peptide-scale alignment: composition-bias correction was disabled (*--comp-bias-corr 0*), the *e-*value cap was set to 1000 (*-e* 1000), and the k-mer prefilter was bypassed (*--prefilter-mode*– 2) so that short peptides (<20 residues) were not dropped during prefiltering. Self-alignments used the default prefilter together with *--add-self-matche*s and *--max-seqs* 1. For prototype-similarity reporting in analog generation, *t* is the prototype; for novelty reporting, *t* ranges over a reference database and the maximum bitscore across the database is taken. A threshold of similarity ≥ 0.60 was adopted throughout this work as the operational definition of analog identity; below this threshold a generated peptide is regarded as a *de novo* design rather than an analog of the prototype.

### Analog generation parameter sweep

The activity-similarity trade-off of analog generation as a function of the exploration strength τ and the property relaxation σ was characterized on two reference sets sampled from DBAASP: 500 active prototypes (MIC ≤ 32 μg/mL against at least one bacterial strain) and 500 inactive prototypes (MIC ≥ 128 μg/mL against all reported strains), provided in Supplementary Data S1. For each prototype, 10 analogs were generated per (τ, σ) configuration across τ ∈ {0.00, 0.05, 0.10, 0.15, 0.20, 0.30, 0.50, 0.70}, σ ∈ {0.00, 0.25, 0.50, 0.75, 1.00}. Per-prototype reporting used the best analog by lowest predicted MIC_APEX_, with similarity to the prototype computed as described above.

### Analog and *de novo* benchmark against HydrAMP

OmegAMP analog modes (AP, AT, AU, AM, AMT) were compared to HydrAMP-A, and OmegAMP *de novo* modes (DU, DT, DP) to HydrAMP-D, using the 500 active and 500 inactive prototypes described above. Targeted modes (AT, DT, and AMT) drew conditioning targets from the therapeutically relevant ranges: charge 2–10, length 5–30 residues, and hydrophobicity –0.5 to +0.8. For HydrAMP-A, the creativity parameter τ ∈ {1, 2.5, 5} was evaluated, and τ = 2.5 was used as the primary comparison, since HydrAMP-A reached its lowest median MIC_APEX_ on active prototypes at this setting. For analog modes, analogs were generated from each prototype, and the one with the lowest MIC_APEX_ was taken as that prototype’s best candidate. For *de novo* modes, generation produced a batch with no per-candidate selection; the prototype set defined the conditioning targets for DP at relaxation σ = 0, fixing charge, length, and hydrophobicity to the prototype values, and was not used for DT or DU. Method-level distributions of MIC_APEX_, best-candidate values for analog modes and all generated sequences for *de novo* modes, were compared by two-sided Mann-Whitney U test against HydrAMP-A (τ = 2.5) for OmegAMP-A modes and against HydrAMP-D for OmegAMP-D modes. The number of valid candidates returned per method, out of the requested 500 (analog) or 50,000 (*de novo*), was reported alongside.

### Inactive prototype selection

Six experimentally inactive peptides were selected as prototypes for analog generation: Mammutin-1^22^, BoCo1^28^, DeNo1047^30^, As-CATH4-6L^27^, OP-145-TII4^11^ and GQ20^29^. Each is inactive against all strains reported in DBAASP (MIC ≥ 128 μg/mL). OP-145-TII4 had itself been generated by HydrAMP as an analog of OP-145^65^ but failed to reach the intended activity.

### Motif derivation for motif-guided generation

Five sequence motifs were used as templates for OmegAMP-AMT and OmegAMP-MT generation, four targeting lipopolysaccharide (LPS) and one targeting DNA. Motifs were encoded as templates of fixed positions (amino acid letters at the residue positions to be preserved) and variable positions (underscores at all other positions); template length defines the length of the generated peptide.

Cecropin-P1 (DBAASPR_569; UniProt P14661): a 31-residue α-helical AMP from *Ascaris suum*. The LPS-binding architecture was taken from the solution NMR structure of cecropin-P1 bound to LPS^31^, in which a cationic KK pair followed by two periodically-spaced isoleucines anchors the amphipathic-helix face engaging lipid A acyl chains. The motif template was “KK <u>I I</u>” (KK at positions 15–16; I at positions 22 and 26).

Sarcotoxin-1A (DBAASPR_1855; UniProt P08375): a 39-residue cecropin-family AMP from *Sarcophaga peregrina*. The N-terminal aromatic-cationic anchor (W followed by paired lysines) was taken from the stable-isotope-assisted NMR characterization of the sarcotoxin-1A–lipid A interaction^35^. The motif template was “_W_KK” (W at position 2; KK at positions 4–5).

Pa4 from pardaxin (DBAASPR_5627): a 33-residue antimicrobial peptide derived from the pardaxin scaffold of *Pardachirus marmoratus*, combining an N-terminal aromatic GFF anchor with a central KP turn and a C-terminal cationic-hydrophobic K_LL_A motif. The LPS-bound conformation was taken from the NMR structure of pardaxin in LPS micelles^34^. The motif template was “ KII P K_LL_A” (KII at positions 8–10; P at position 13; K at position 16; LL at positions 18–19; A at position 21).

LG21 (temporin LG21; DBAASPS_7148): a 21-residue synthetic peptide with an LPS-binding C-terminal extension^32,33^. The motif template was “ GWKRKRFG” (GWKRKRFG at positions 14–21).

bZIP from GCN4 (PDB 1GD2): a 30-residue leucine-zipper peptide from the basic-region leucine-zipper transcription factor GCN4 of *Saccharomyces cerevisiae*^36,38^. The dimerization core was taken from the GCN4 coiled-coil structure, in which leucines repeat every seventh residue to form the zipper. The motif template was “ L V L N L V L V_” (L at positions 4, 11, 18 and 25; V at 8, 22 and 29; N at 15).

### Wet-lab candidate selection

Each pool was then filtered and ranked by the same procedure. Duplicate sequences and sequences identical to a known AMP were removed, and the synthesis-feasibility filter (**Synthesis filtering** section) was applied to every pool. Analog and motif-guided candidates were additionally required to retain at least 60% similarity to their prototype (computed as in the **Sequence similarity** section), and motif-guided candidates to preserve the template residues. The bZIP designs were exempt from the similarity requirement, since the bZIP parent is not antimicrobial and similarity to it is not a design constraint. Surviving candidates were ranked in lexicographic order by the OmegAMP general-classifier call, then the APEX rank (number of the 34 APEX strain classifiers predicting MIC ≤ 32 μM), then the OmegAMP rank (number of the 12 strain– and species-specific OmegAMP classifiers positive at probability > 0.5). The OmegAMP general-classifier and the 12 strain– and species-specific OmegAMP classifiers are auxiliary discriminative models trained alongside the generative framework on the same labeled data, each returning the probability that a sequence is antimicrobial, overall or against the corresponding strain or species (full specification in the OmegAMP paper^44^). From each pool, up to 500 top-ranked candidates were retained, and the final candidates advanced to synthesis were chosen from these by expert assessment, giving 95 de novo (85 Ω-DP, 10 Ω-DT), 59 analog (22 Ω-AP, 29 Ω-AT, 8 Ω-AU) and 50 motif-guided (10 per template) peptides. With the 11 reference prototypes, the full wet-lab library comprised 215 peptides, with polymyxin B and levofloxacin as antibiotic controls.

### Synthesis filtering

Generated peptides were filtered by a set of synthesis-feasibility rules implemented in the OmegAMP repository. A peptide was retained only if it met all of the following criteria: hydrophilic residue abundance ∈ [30%, 70%], where hydrophilic residues are K, R, H, D, E, S, T, N, Q, Y; no run of more than two consecutive glycines; no occurrence of any of 24 aggregation-prone motifs (compiled from expert knowledge; including VLVL, WWYF, KLLL, STVIIE, VQIVYK, KLVFFA, LVFFAEDVGSNK, NFGAIL, DFNKF and related sequences); proline content ≤ 20% of residues; at most one cysteine (avoiding intermolecular disulfide bonds during synthesis); and net charge at physiological pH ∈ [2, 10]. Filtering was applied after generation and before candidate ranking.

### Peptide synthesis

Peptides were obtained from AAPPTec (Louisville, KY) and synthesized by solid-phase peptide synthesis using the Fmoc strategy.

### Minimum inhibitory concentration determination

MICs were determined by broth microdilution. Peptides were dispensed into untreated polystyrene 96-well microtiter plates and two-fold serially diluted in sterile water to final concentrations ranging from 0.03125 to 64 μmol L⁻¹. Bacterial suspensions were prepared in LB medium at 4 × 10⁶ CFU mL⁻¹ and mixed 1:1 with the peptide dilutions. Plates were incubated at 37 °C for 24 h, and the MIC was defined as the lowest peptide concentration that completely inhibited visible bacterial growth. All assays were performed in three independent replicates.

## Hemolysis assay

Hemolytic activity was evaluated by measuring hemoglobin release from human red blood cells (RBCs). Human RBCs from a male A-donor were obtained from Zen-Bio Inc. (Durham, North Carolina, USA) as heparin-anticoagulated blood. RBCs were washed four times with PBS (pH 7.4) by centrifugation at 800 × g for 10 min. Washed RBCs were diluted 200-fold in PBS, and 75 μL of the cell suspension was mixed with 75 μL of peptide solution, giving final peptide concentrations ranging from 2 to 128 μmol L⁻¹. Samples were incubated for 4 h at room temperature and then centrifuged at 1,300 × g for 10 min to pellet intact cells and debris. Supernatants (100 μL) were transferred to a new 96-well plate, and absorbance was measured at 405 nm using a plate reader. Percent hemolysis was calculated by normalizing each sample to PBS-treated RBCs as the negative control and 1% (v/v) SDS in PBS as the positive control.

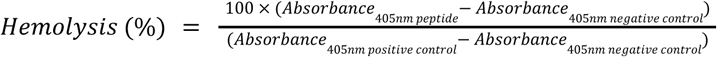

## Human embryonic kidney 293T cells cytotoxicity assay

HEK293T cell cytotoxicity was assessed using the 3-(4,5-dimethylthiazol-2-yl)-2,5-diphenyltetrazolium bromide (MTT) assay. Cells were cultured in high-glucose Dulbecco’s modified Eagle’s medium (DMEM) supplemented with 10% fetal bovine serum and 1% penicillin-streptomycin and maintained at 37 °C in a humidified atmosphere containing 5% CO₂. One day before treatment, HEK293T cells were seeded into 96-well plates at 10,000 cells per well in 100 μL of medium and allowed to attach overnight. Cells were then treated with peptides at concentrations ranging from 8 to 128 μmol L⁻¹ and incubated for 24 h.

After treatment, peptide-containing medium was removed and replaced with 100 μL of MTT reagent prepared at 0.5 mg mL⁻¹ in phenol red-free medium. Plates were incubated for 4 h at 37 °C under 5% CO₂ to allow formazan crystal formation. The resulting crystals were dissolved in 0.04 mol L⁻¹ hydrochloric acid in anhydrous isopropanol, and absorbance was measured at 570 nm using a plate reader. Cell viability was calculated relative to untreated controls. All experiments were performed in three biological replicates.

## Circular dichroism spectroscopy

Circular dichroism (CD) spectroscopy was performed using a J-1500 circular dichroism spectropolarimeter (Jasco) at the Biological Chemistry Resource Center, University of Pennsylvania. Spectra were acquired at 25 °C using a quartz cuvette with a 1.0 mm path length. For each peptide, spectra were recorded from 260 to 190 nm at a scan rate of 50 nm min⁻¹, with a bandwidth of 0.5 nm, and averaged over three accumulations. Peptides were measured at 50 μmol L⁻¹ in water, 60% trifluoroethanol (TFE) in water, 50% methanol (MeOH) in water, or 10 mmol L⁻¹ sodium dodecyl sulfate (SDS) in water. Matching solvent baselines were collected before peptide measurements and subtracted from the corresponding spectra. A Fourier transform filter was applied to reduce background noise. Secondary-structure fractions were estimated from the processed spectra using the single-spectrum analysis tool on the BeStSel server.

## Outer-membrane permeabilization assay

Outer-membrane permeabilization was evaluated using the N-phenyl-1-naphthylamine (NPN) uptake assay. *A. baumannii* ATCC 19606 was grown to an OD₆₀₀ of 0.4, harvested by centrifugation at 9,391 × g for 3 min, washed, and resuspended in 5 mmol L⁻¹ HEPES buffer (pH 7.4) containing 5 mmol L⁻¹ glucose. The bacterial suspension was added to white 96-well plates at 100 μL per well, followed by 4 μL of NPN solution at 0.5 mmol L⁻¹. Peptides diluted in sterile water were then added to each well, and fluorescence was monitored over 45 min using excitation and emission wavelengths of 350 and 420 nm, respectively. At each time point, fluorescence was normalized to the untreated control containing buffer, bacteria, and NPN. Replicate fluorescence values were first averaged, and the percentage difference relative to the untreated control was calculated as:

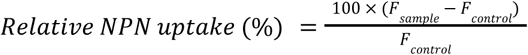

Where *F_sample_* and *F_control_* represent the mean fluorescence intensities of the peptide-treated and untreated samples, respectively, at the corresponding time point.

The resulting relative-fluorescence time courses were fitted in GraphPad Prism using an unweighted least-squares third-order polynomial model,

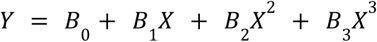

without outlier exclusion. MaxRel was defined as the maximum relative fluorescence value predicted by the fitted curve within the experimental time interval. The area under the fitted relative-fluorescence curve (AUC) was calculated over the complete recording interval using Y = 0 as the baseline. AUC values are therefore expressed in relative-fluorescence percentage × time.

## Membrane depolarization assay

Cytoplasmic membrane depolarization was evaluated using the membrane potential-sensitive dye 3,3′-dipropylthiadicarbocyanine iodide [DiSC₃(5)]. *A. baumannii* ATCC 19606 was grown to mid-logarithmic phase, harvested by centrifugation at 9,391 × g for 3 min, washed, and resuspended to an OD₆₀₀ of 0.05 in HEPES buffer (pH 7.2) containing 20 mmol L⁻¹ glucose and 0.1 mol L⁻¹ KCl. DiSC₃(5) was added to the bacterial suspension at 20 μmol L⁻¹, and 100 μL of the mixture was added to each well of a 96-well plate. The suspension was incubated for 15 min to allow dye incorporation into the bacterial membrane and stabilization of fluorescence. Peptides diluted in sterile water were then mixed 1:1 with the bacterial suspension to final concentrations corresponding to their MIC values. Membrane depolarization was monitored by measuring fluorescence over 60 min using excitation and emission wavelengths of 622 and 670 nm, respectively.

At each time point, fluorescence was normalized to the untreated control containing buffer, bacteria, and DiSC_3_(5). Replicate fluorescence values were first averaged, and the percentage difference relative to the untreated control was calculated as:

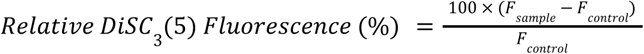

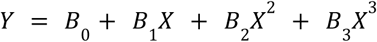

## LPS engagement (BC displacement) assay

Peptide engagement with lipopolysaccharide (LPS) was evaluated using a BODIPY-TR-cadaverine (BC) displacement assay. BC associates with LPS, resulting in a lower fluorescence signal in the LPS-bound state. Peptides that interact with LPS compete with BC and promote its displacement, leading to fluorescence recovery. Thus, increased BC displacement indicates greater relative competition for LPS binding under the assay conditions.

BC was incubated with LPS in assay buffer to establish the LPS-bound reference state. Peptides diluted in sterile water were then added to the LPS–BC mixture, and fluorescence was measured using a microplate reader. Polymyxin B, a known LPS-binding molecule, was included as a reference control but was not used to define the upper normalization limit. To assess whether peptide–LPS engagement was maintained in the presence of divalent cations, the assay was also performed under a calcium-challenge condition by including CaCl₂ in the assay buffer.

BC displacement was normalized independently for each experimental condition. The fluorescence of BC incubated with LPS in the absence of peptide was defined as 0% displacement, whereas the fluorescence of free BC in buffer without LPS was defined as 100% displacement. BC displacement was calculated as:

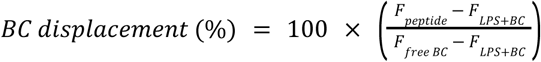

where *F_peptide_* is the fluorescence measured after addition of the peptide to the LPS–BC mixture, *F_LPS+BC_* is the fluorescence of BC bound to LPS in the absence of peptide, and *F_free BC_* is the fluorescence of BC in buffer without LPS. For calcium-challenge experiments, both reference signals were measured in the presence of the same CaCl_2_ concentration used for the corresponding peptide samples. Higher displacement values indicate greater relative competition with BC for interaction with LPS but should not be interpreted as an absolute measurement of LPS-binding affinity.

## DNA perturbation by circular dichroism assay

Peptide-induced perturbation of DNA was evaluated by circular dichroism (CD) spectroscopy using calf thymus DNA. DNA was prepared in assay buffer and incubated either alone or with peptide before spectral acquisition. CD spectra were collected under the same instrumental conditions described for peptide secondary-structure measurements. The spectrum of DNA alone was used as the reference, and each peptide–DNA spectrum was compared with this control to assess changes in the DNA CD signature.

Analysis focused on the two characteristic DNA spectral features: the negative band near 245 nm and the positive band near 275 nm. Peptide-induced changes were quantified using two complementary parameters: amplitude shift and wavelength shift. For each band, the wavelength of the corresponding maximum or minimum in the DNA-only spectrum was used as the reference wavelength. The amplitude shift was calculated as:

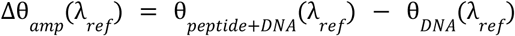

where θ*_peptide+DNA_* and θ*_DNA_* are the CD signals measured for the peptide-DNA and DNA-only conditions, respectively, and λ*_ref_* is the wavelength of the corresponding feature in the DNA-only spectrum.

For the positive band near 275 nm, a positive amplitude shift indicates an increase in the positive CD signal, whereas a negative value indicates a decrease. For the negative band near 245 nm, a positive shift indicates that the signal became less negative, whereas a negative shift indicates that it became more negative.

The wavelength shift was determined by comparing the position of the corresponding spectral extremum in the peptide–DNA and DNA-only spectra. For the positive band, the shift was calculated as:

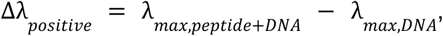

whereas the shift of the negative band was calculated as:

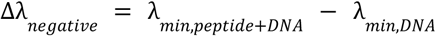

Positive values indicate a shift toward longer wavelengths, whereas negative values indicate a shift toward shorter wavelengths. Amplitude and wavelength shifts were interpreted together as comparative measures of peptide-induced changes in the DNA CD spectrum. These parameters indicate changes consistent with peptide–DNA interaction or DNA structural perturbation but were not interpreted as direct measurements of binding affinity. At higher peptide concentrations, spectral changes may reflect a combination of DNA association, conformational perturbation, and formation of higher-order peptide–DNA assemblies.

## Resistance to proteolytic degradation assay

Peptide stability against enzymatic degradation was evaluated by incubating the peptides in an aqueous solution containing 25% human serum. Peptides at a concentration of 10 mg ml^-1^ were incubated for 6 h with 25% human serum in water (Zen-Bio; healthy donor, blood type A^-^). Aliquots were collected at 0, 0.5, 1, 2, 4 and 6 h. Immediately after collection, each aliquot was mixed with 10 μl of trifluoroacetic acid and incubated for 10 min to terminate enzymatic activity.

Samples were analyzed using a Waters Acquity ultrahigh-performance liquid chromatography-mass spectrometry system equipped with a photodiode array detector, with data collected at 220 nm, and a Waters single quadrupole detector 2. Chromatographic separation was performed using a Waters XBridge C_18_ column (3.5 µm, 4.6 mm × 50 mm). The mobile phases consisted of 100% water supplemented with 0.1% (v/v) formic acid as solvent A and 100% acetonitrile as solvent B. Both solvents were Fisher Optima grade. An injection volume of 50 μl was used. Ionization was conducted in both positive and negative electrospray ionization modes, with a mass-scanning range of m/z 100-3,000. The chromatographic gradient increased solvent B from 5% to 95% over 5 min.

The percentage of intact peptide remaining at each time point was determined by integrating the area under the peptide peak and normalizing it to the corresponding peak area measured at the initial time point (t = 0). All experiments were performed in triplicate.

## Quantification and Statistical Analysis

Dose-response curves for hemolysis and HEK293T cytotoxicity were fitted by non-linear sigmoidal regression in GraphPad Prism v.10 to obtain HC_50_ and CC_50_. NPN and DiSC_3_(5) time courses were fitted by non-linear regression to obtain MaxRel and AUC summaries. Downstream statistical analysis was performed in Python with *scipy.stats*, *statsmodels.stats.multitest*, and *pingouin*. Cross-group comparisons across more than two groups (e.g., across prototype families) used the Kruskal-Wallis test (*pingouin.kruskal*), with the effect size reported as η² = (*H* – *k* + 1)/(*n* – *k*), where *H* is the Kruskal-Wallis statistic, *k* the number of groups, and *n* the total number of observations. Pairwise comparisons used the two-sided Mann-Whitney U test (*pingouin.mwu* or *scipy.stats.mannwhitneyu* with *alternative=’two-sided’*), with the rank-biserial correlation *r* as the effect size. For the murine efficacy experiments (**Figure 6**), bacterial burden (log₁₀ CFU per gram) in each day-block was first assessed by a Kruskal-Wallis test across all groups, and each treatment group was then compared with the untreated control by a two-sided exact Mann-Whitney U test (*scipy.stats.mannwhitneyu* with method=’*exact*’), without multiple-comparison correction. Univariate correlations between continuous variables (e.g., potency versus safety, sequence novelty versus activity, physicochemical properties versus MIC) were assessed by Spearman rank correlation (*scipy.stats.spearmanr*). Partial Spearman correlations controlling for covariates (e.g., charge controlled for hydrophobicity and length) were computed with *pingouin.partial_corr* using *method=’spearman’*. Linear regression trend lines were fitted with *scipy.stats.linregress*. The number of biological replicates (*n*), the test used, the test statistic, and the exact *p*-value are reported in the corresponding figure legend. Significance thresholds are denoted as * *p* < 0.05, ** *p* < 0.01, *** *p* < 0.001; *n.s.*, not significant. Sample sizes were chosen to match the standards in the field for *in vitro* antimicrobial characterization and were not predetermined by power analysis. Investigators performing wet-lab assays were blinded to peptide design details (generation mode, prototype family, and sequence); each peptide was identified only by an anonymized ID during assay execution and data acquisition.

## Additional Resources

OmegAMP dashboard for interactive exploration of the designed peptides and their experimental characterization: https://szczurek-lab.github.io/omegamp-dashboard/.

## Extended Data Figures

**Extended Data Fig. 1.**
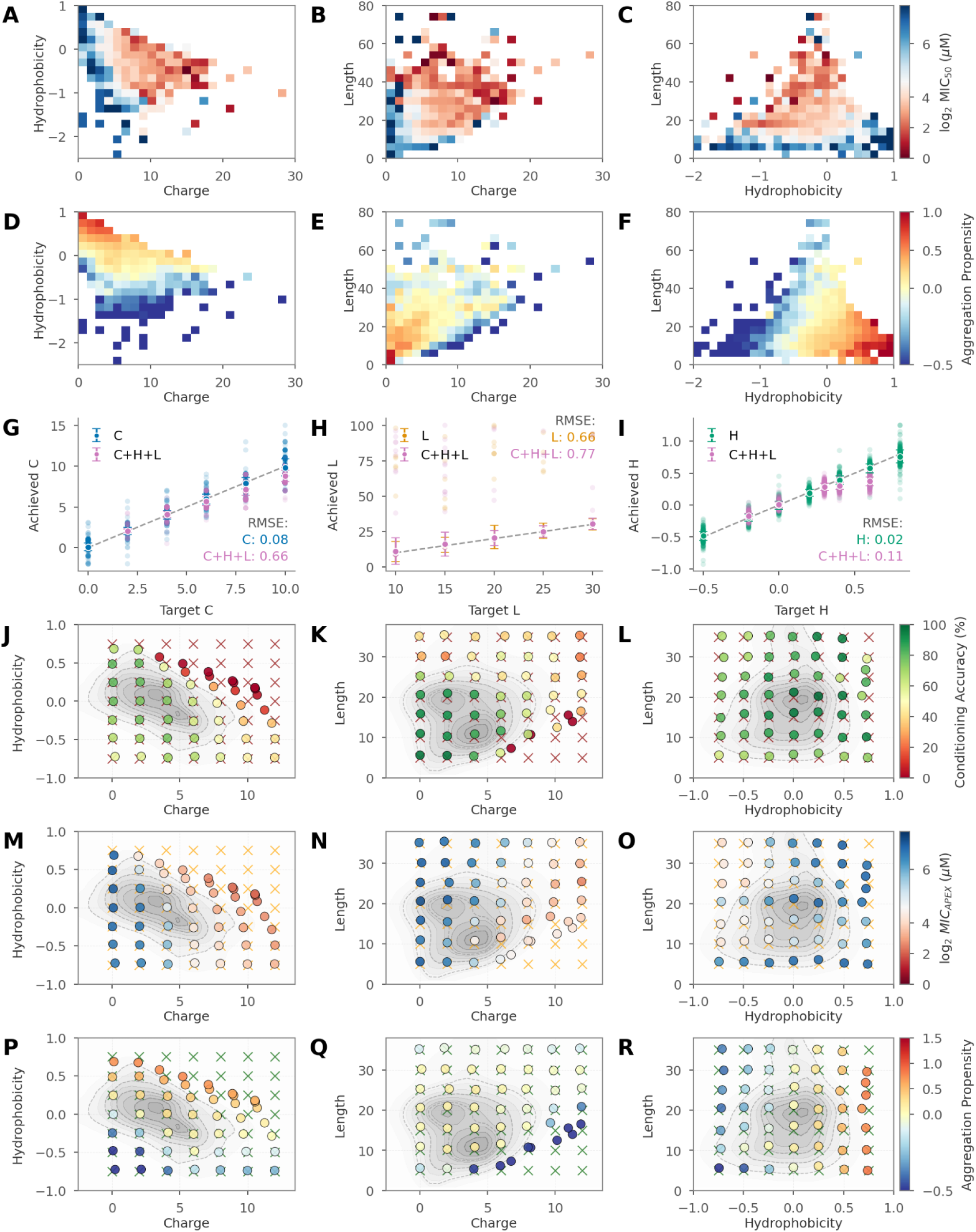
Physicochemical constraints define the boundaries of targeted generation. (**A-C**) Experimental MIC landscape across physicochemical space. MIC₅₀ (log₂ μM) from *n = 5,207* experimentally characterized peptides, computed as the median across species and binned by charge-hydrophobicity (**A**), charge-length (**B**), and hydrophobicity-length (**C**). Lower values (red) indicate higher antimicrobial potency. (**D-F**) Experimental aggregation propensity across the same peptides and projections as (A–C): charge-hydrophobicity (**D**), charge-length (**E**), and hydrophobicity-length (**F**). Higher values (red) indicate greater aggregation tendency. (**G**-**I**) Achieved versus target values under single-property conditioning (colored points; 500 peptides per target) and simultaneous three-property conditioning (pink; C+L+H coupled targets, see main text), for charge (**G**), length (**H**), and hydrophobicity (**I**). Dashed line, perfect conditioning. Root-mean-square error (RMSE) is reported for each property under both regimes. (**J**-**L**) Conditioning accuracy, the fraction of generated peptides within ±0.25 charge units, ±0.1 hydrophobicity units, and exact length match, across 49 grid coordinates per property pair: charge-hydrophobicity (**J**), charge-length (**K**), and hydrophobicity-length (**L**). Gray contours, training data density; crosses, target coordinates. (**M**-**O)** Predicted activity (log₂ MIC_APEX_, μM) for generated peptides at each grid coordinate, across the same projections as (J–L): charge-hydrophobicity (**M**), charge-length (**N**), and hydrophobicity-length (**O**). Lower values (red) indicate higher predicted antimicrobial potency. (**P**-**R**) Predicted aggregation propensity for generated peptides at each grid coordinate, across the same projections as (J–L): charge-hydrophobicity (**P**), charge-length (**Q**), and hydrophobicity-length (**R**). Higher values (red) indicate greater predicted aggregation tendency.

**Extended Data Fig. 2.**
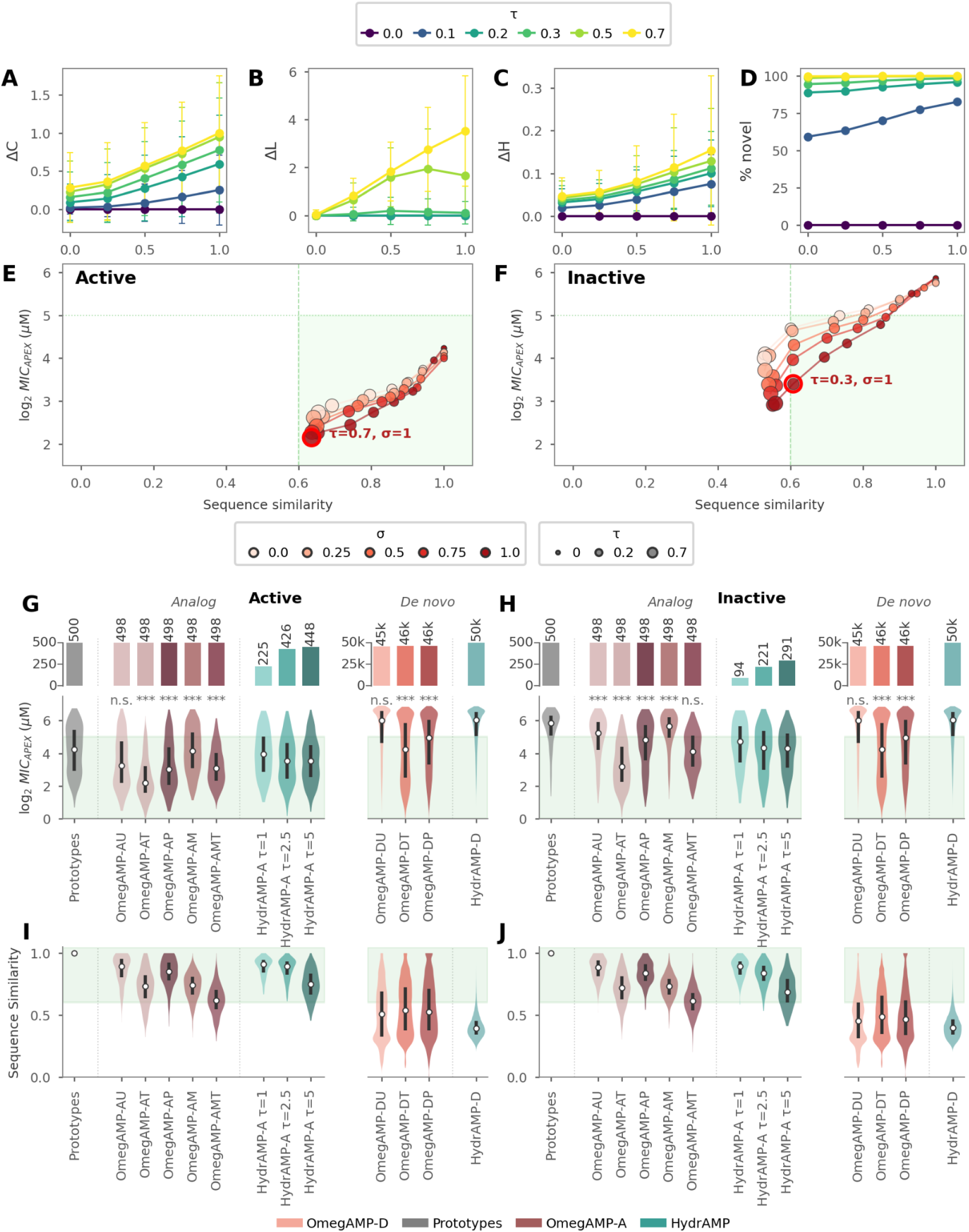
Controlled analog generation balances activity and sequence preservation. (**A**–**C**) Property deviation Δ (mean absolute error, MAE) between analogs and prototypes as a function of property relaxation (σ), colored by exploration strength (τ), for net charge (ΔC, **A**), length (ΔL, **B**), and mean hydrophobicity (Δ<H>, **C**). Error bars denote standard deviation. (**D**) Percentage of novel analogs as a function of σ, colored by τ. (**E**) Activity-similarity trade-off across the τ-σ parameter space for active prototypes. Color encodes σ; point size encodes τ. Green shading marks the desirable regime (≥60% sequence similarity, ≤32 μM MIC_APEX_). The labeled point indicates the optimum (*τ = 0.7, σ = 1.0*) (see also **Extended Data Fig. 3A and 3B**). (**F**) As in (E) for inactive prototypes; the labeled point indicates the optimum (τ = 0.3, σ = 1.0) (see also **Extended Data Fig. 3C and 3D**). (**G**, **H**) Benchmarking OmegAMP analog and de novo modes against HydrAMP for active (**G**) and inactive (**H**) prototypes. Top bars, number of valid candidates returned per method (annotated above each bar; target 500 analog, 50,000 de novo). Violins, log₂ MIC_APEX_ of the best candidate per prototype. Gray, prototype distributions; white points, medians; bars, interquartile ranges; green shading, MIC_APEX_ ≤ 32 μM. Asterisks, Mann-Whitney U (MW; two-sided) versus HydrAMP-A τ = 2.5 for OmegAMP-A modes and versus HydrAMP-D for OmegAMP-D modes: *\* – p < 0.05, ** – p < 0.01, *** – p < 0.001, n.s.* – not significant. (**I**, **J**) Sequence similarity of the best candidate per prototype (maximum normalized local-alignment bitscore, AMP-derived substitution matrix; see Methods) for active (**I**) and inactive (**J**) prototypes. Green shading, similarity ≥ 0.6.

**Extended Data Fig. 3.**
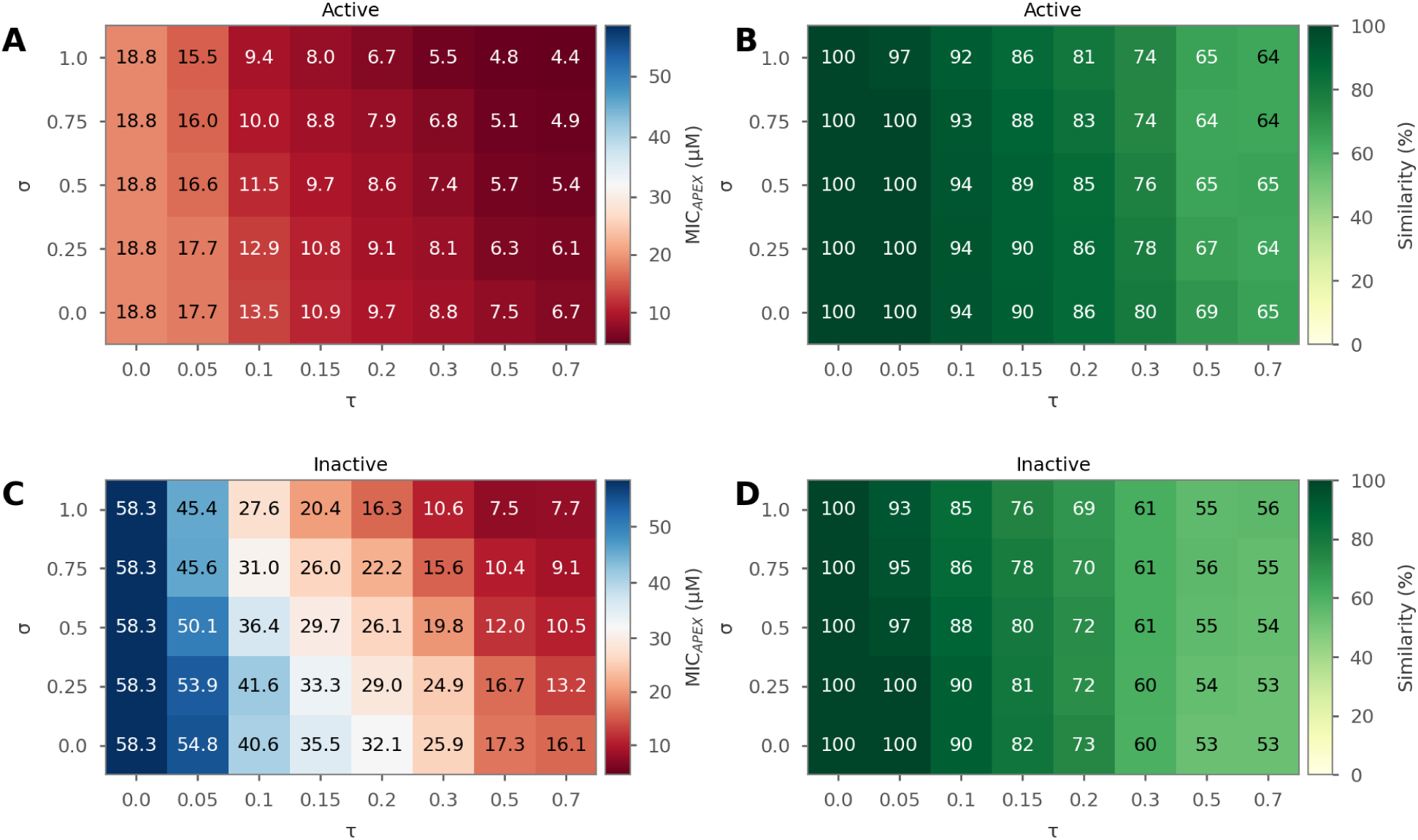
Detailed parameter sweeps for analog generation, related to Extended Data Fig. 2. (**A**, **B**) Median predicted MIC (MIC_APEX_, μM; A) and median sequence similarity (%; B) across the τ-σ parameter space for the best analog per active prototype, selected by predicted activity. (**C**, **D**) Median predicted MIC (C) and median sequence similarity (D) across the τ-σ parameter space for the best analog per inactive prototype.

**Extended Data Fig. 4.**
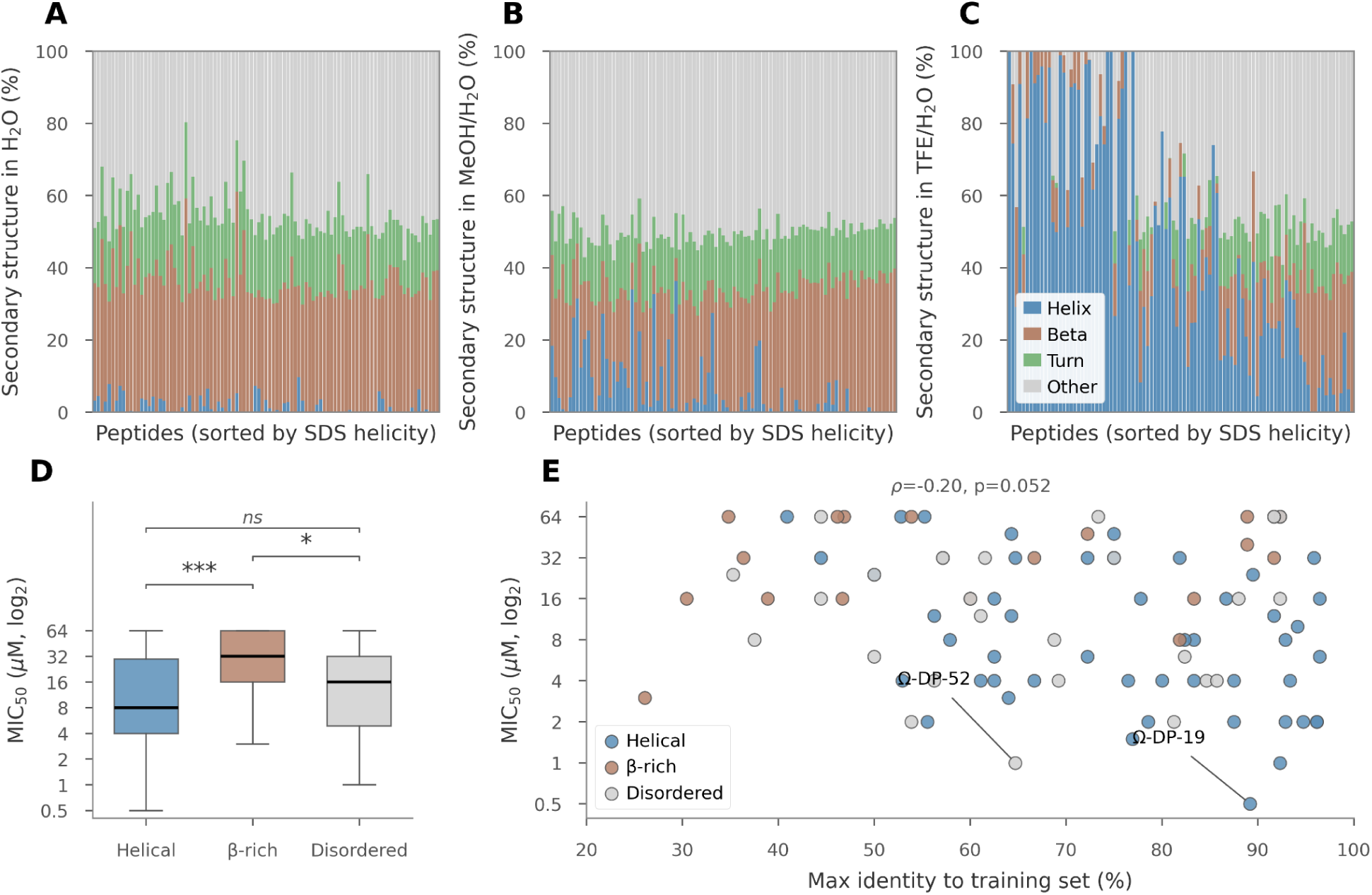
Cross-medium secondary structure, structural-class activity, and sequence novelty of *de novo* peptides, related to Figure 3. (**A**) Secondary structure composition of *de novo* peptides in H₂O determined by CD. Bars represent fractional composition of α-helix, β-strand, turn, and other structures, sorted by decreasing SDS helicity (same peptide ordering as **Figure 3B**). (**B**) Secondary structure composition in MeOH/H₂O, sorted as in (A). (**C**) Secondary structure composition in TFE/H₂O, sorted as in (A). (**D**) MIC₅₀ by secondary structure class in SDS. Helical: α-helix > 50%; β-rich: α-helix ≤ 50% and β-strand > 20%; disordered: α-helix ≤ 50% and β-strand ≤ 20%. Brackets indicate pairwise MW tests with significance encoded as *\*\*\* p < 0.001, ** p < 0.01, * p < 0.05, ns p ≥ 0.05.* (**E**) Sequence novelty versus antimicrobial activity. Best gapless, length-matched identity to the training-set AMPs plotted against MIC₅₀ across the 20-strain panel. Points are colored by secondary structure class in SDS as in (D). The Spearman rank correlation coefficient ρ and its *p-*value are shown. Leads Ω-DP-19 and Ω-DP-52 are annotated.

**Extended Data Fig. 5.**
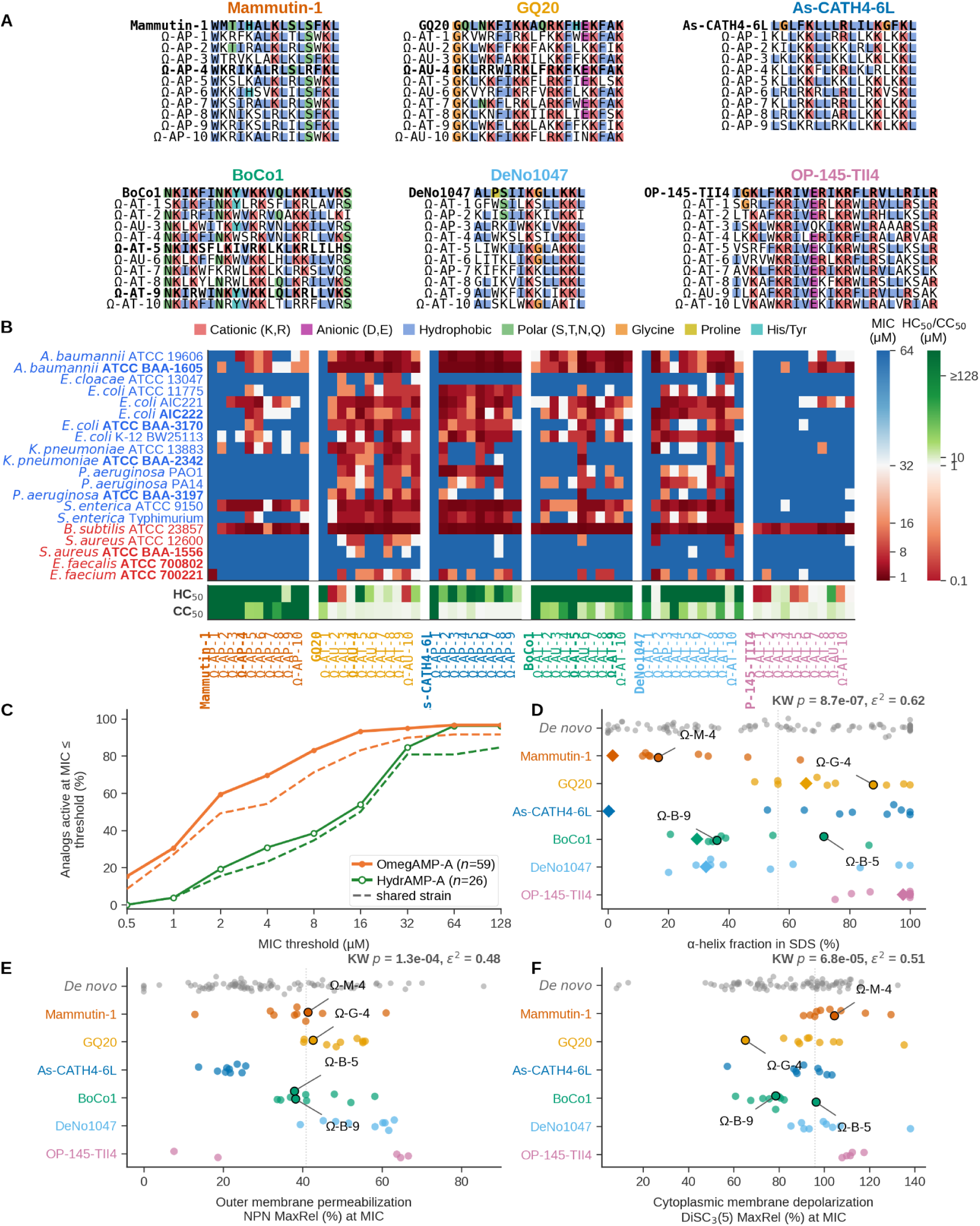
Wet-lab characterization of analogs from inactive prototypes, related to Figure 4. (**A**) Sequence alignments for the six inactive-prototype families. Prototype on top, analogs below; analog suffix indicates conditioning (AP, prototype-derived; AT, property-targeted; AU, unconditional). Residues colored by physicochemical class (legend); **Table 1** leads in bold. (**B**) MIC heatmap for prototypes and the 59 analogs across the 20-strain panel; red, potent; blue, resistant. HC₅₀ and CC₅₀ shown below (green, favorable; red, toxic). Gram-negative strains, blue labels; Gram-positive, red; MDR isolates, bold. (**C**) Fraction of analogs with MIC ≤ threshold against at least one of the four species assayed in both studies (solid) and against *A. baumannii* ATCC BAA-1605, the only strain assayed in both (dashed), for OmegAMP-A (orange, *n* = 59) and HydrAMP-A (green, *n* = 26). OmegAMP-A strains: *E. coli* ATCC 11775, AIC221, AIC222, ATCC BAA-3170 and K-12 BW25113; *S. aureus* ATCC 12600 and ATCC BAA-1556; *A. baumannii* ATCC 19606 and ATCC BAA-1605; *P. aeruginosa* PAO1, PA14 and ATCC BAA-3197. HydrAMP-A strains: *E. coli* ATCC 25922 and ATCC 43827; *S. aureus* ATCC 33591; *A. baumannii* ATCC BAA-1605; *P. aeruginosa* ATCC 9027. HydrAMP-A MICs converted from µg/mL to µM per peptide molecular weight. (**D**) α-Helix fraction in SDS micelles by prototype family; *de novo* set as reference (top). Diamonds, prototypes; leads outlined in black. Dashed line, 56% threshold for predominantly α-helical structure. KW *p* and ε² report variance attributable to the prototype family. (**E**, **F**) Outer membrane permeabilization (NPN MaxRel, %, **E**) and cytoplasmic membrane depolarization (DiSC₃(5) MaxRel, %, **F**) by prototype family, each measured at each analog’s MIC. Diamonds, prototypes; dashed line, global median. KW p and ε² shown.

**Extended Data Fig. 6.**
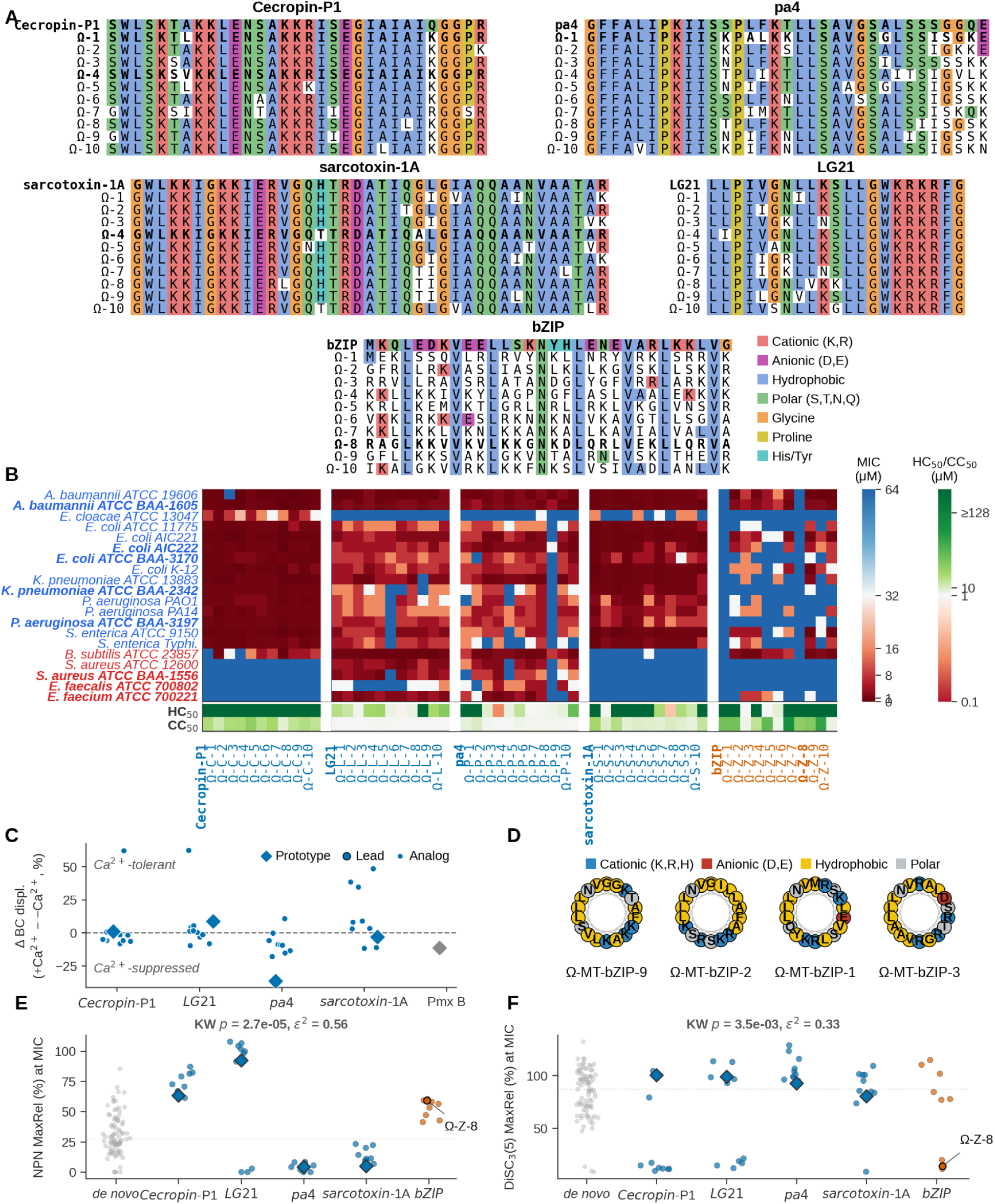
Wet-lab characterization of motif-guided analogs, related to Figure 5. **(A)** Sequence alignments for the four LPS-engaging families (cecropin-P1, pa4, sarcotoxin-1A, LG21) and the DNA-engaging family (bZIP). Prototype on top in each panel, analogs Ω-1 through Ω-10 below. Residues colored by physicochemical class (legend); **Table 1** leads in bold. **(B)** MIC heatmap for prototypes and the 50 motif-guided designs across the 20-strain panel; potent: red, weak: blue. HC₅₀ and CC₅₀ shown below (green, favorable; red, toxic). Gram-negative strains, blue labels; Gram-positive, red; MDR isolates, bold. **(C)** Calcium dependence of BC displacement for the four LPS-engaging families and the polymyxin B reference. ΔBC displacement (%) is the difference in BODIPY-cadaverine displacement at 32 μM peptide between +Ca²⁺ and –Ca²⁺ conditions (+Ca²⁺ minus –Ca²⁺). Positive ΔBC, Ca²⁺-tolerant or Ca²⁺-enhanced; negative ΔBC, Ca²⁺-suppressed. Dashed line, zero. Diamonds, prototypes and polymyxin B (Pmx B, gray); circles outlined in black, leads (Ω-C-1, Ω-P-1, Ω-S-4); other circles, analogs. **(D)** Helical wheels of four bZIP designs ordered by decreasing DNA perturbation strength. Residues 1–18 projected onto a single wheel (100° per residue), colored by physicochemical class. **(E)** Outer membrane permeabilization (NPN MaxRel, %) at MIC by prototype family; *de novo* set as reference (left). Diamonds, prototypes; leads outlined in black and labeled. Dashed line, global median across the wet-tested set. KW *p* and ε² report variance attributable to the prototype family. **(F)** Cytoplasmic membrane depolarization [DiSC₃(5) MaxRel, %] at MIC by prototype family. Conventions as in (E).

## References

1. Murray, C.J.L., Ikuta, K.S., Sharara, F., Swetschinski, L., Aguilar, G.R., Gray, A., Han, C., Bisignano, C., Rao, P., Wool, E., et al. (2022). Global burden of bacterial antimicrobial resistance in 2019: a systematic analysis. The Lancet 399, 629–655. 10.1016/S0140-6736(21)02724-0.

2. Lazarus, G., Caddey, B., Dean, A., Wangsaputra, V.K., Febrina, F., Radiani, S.P., Xiong, Y., Conly, J.M., Barkema, H.W., Kleef, E. van, et al. (2026). Antimicrobial resistance in bacterial pathogens causing community-acquired and hospital-acquired lower respiratory tract infections (2010–24): a systematic review and meta-analysis. Lancet Microbe 7. 10.1016/j.lanmic.2026.101361.

3. Antibacterial pipeline trends and recommendations to enhance research and development (2024). (World Health Organization).

4. Melchiorri, D., Rocke, T., Alm, R.A., Cameron, A.M., and Gigante, V. (2025). Addressing urgent priorities in antibiotic development: insights from WHO 2023 antibacterial clinical pipeline analyses. Lancet Microbe 6, 100992. 10.1016/j.lanmic.2024.100992.

5. Fjell, C.D., Hiss, J.A., Hancock, R.E.W., and Schneider, G. (2012). Designing antimicrobial peptides: form follows function. Nat. Rev. Drug Discov. 11, 37–51. 10.1038/nrd3591.

6. Spohn, R., Daruka, L., Lázár, V., Martins, A., Vidovics, F., Grézal, G., Méhi, O., Kintses, B., Számel, M., Jangir, P.K., et al. (2019). Integrated evolutionary analysis reveals antimicrobial peptides with limited resistance. Nat. Commun. 10, 4538. 10.1038/s41467-019-12364-6.

7. Zasloff, M. (2002). Antimicrobial peptides of multicellular organisms. Nature 415, 389–395. 10.1038/415389a.

8. Mookherjee, N., Anderson, M.A., Haagsman, H.P., and Davidson, D.J. (2020). Antimicrobial host defence peptides: functions and clinical potential. Nat. Rev. Drug Discov. 19, 311–332. 10.1038/s41573-019-0058-8.

9. Brogden, K.A. (2005). Antimicrobial peptides: pore formers or metabolic inhibitors in bacteria? Nat. Rev. Microbiol. 3, 238–250. 10.1038/nrmicro1098.

10. Lazzaro, B.P., Zasloff, M., and Rolff, J. (2020). Antimicrobial peptides: Application informed by evolution. Science 368, eaau5480. 10.1126/science.aau5480.

11. Szymczak, P., Możejko, M., Grzegorzek, T., Jurczak, R., Bauer, M., Neubauer, D., Sikora, K., Michalski, M., Sroka, J., Setny, P., et al. (2023). Discovering highly potent antimicrobial peptides with deep generative model HydrAMP. Nat. Commun. 14, 1453. 10.1038/s41467-023-36994-z.

12. Van Oort, C.M., Ferrell, J.B., Remington, J.M., Wshah, S., and Li, J. (2021). AMPGAN v2: Machine Learning-Guided Design of Antimicrobial Peptides. J. Chem. Inf. Model. 61, 2198–2207. 10.1021/acs.jcim.0c01441.

13. Dean, S.N., Alvarez, J.A.E., Zabetakis, D., Walper, S.A., and Malanoski, A.P. (2021). PepVAE: Variational Autoencoder Framework for Antimicrobial Peptide Generation and Activity Prediction. Front. Microbiol. 12. 10.3389/fmicb.2021.725727.

14. Wang, X.-F., Tang, J.-Y., Sun, J., Dorje, S., Sun, T.-Q., Peng, B., Ji, X.-W., Li, Z., Zhang, X.-E., and Wang, D.-B. (2024). ProT-Diff: A Modularized and Efficient Strategy for De Novo Generation of Antimicrobial Peptide Sequences by Integrating Protein Language and Diffusion Models. Adv. Sci. 11, e2406305. 10.1002/advs.202406305.

15. Szymczak, P., Zarzecki, W., Wang, J., Duan, Y., Wang, J., Coelho, L.P., de la Fuente-Nunez, C., and Szczurek, E. AI-Driven Antimicrobial Peptide Discovery: Mining and Generation. Acc. Chem. Res. 58, 1831–1846. 10.1021/acs.accounts.0c00594.

16. Szymczak, P., and Szczurek, E. (2023). Artificial intelligence-driven antimicrobial peptide discovery. Curr. Opin. Struct. Biol. 83, 102733. 10.1016/j.sbi.2023.102733.

17. Das, P., Sercu, T., Wadhawan, K., Padhi, I., Gehrmann, S., Cipcigan, F., Chenthamarakshan, V., Strobelt, H., dos Santos, C., Chen, P.-Y., et al. (2021). Accelerated antimicrobial discovery via deep generative models and molecular dynamics simulations. Nat. Biomed. Eng. 5, 613–623. 10.1038/s41551-021-00689-x.

18. Torres, M.D.T., Chen, L.T., Wan, F., Chatterjee, P., and de la Fuente-Nunez, C. (2025). Generative latent diffusion language modeling yields anti-infective synthetic peptides. Cell Biomater. 1, 100183. 10.1016/j.celbio.2025.100183.

19. Trippe, B.L., Yim, J., Tischer, D., Baker, D., Broderick, T., Barzilay, R., and Jaakkola, T. (2023). Diffusion probabilistic modeling of protein backbones in 3D for the motif-scaffolding problem. Preprint at arXiv, 10.48550/arXiv.2206.04119 https://doi.org/10.48550/arXiv.2206.04119.

20. Watson, J.L., Juergens, D., Bennett, N.R., Trippe, B.L., Yim, J., Eisenach, H.E., Ahern, W., Borst, A.J., Ragotte, R.J., Milles, L.F., et al. (2023). De novo design of protein structure and function with RFdiffusion. Nature 620, 1089–1100. 10.1038/s41586-023-06415-8.

21. Pirtskhalava, M., Amstrong, A.A., Grigolava, M., Chubinidze, M., Alimbarashvili, E., Vishnepolsky, B., Gabrielian, A., Rosenthal, A., Hurt, D.E., and Tartakovsky, M. (2020). DBAASP v3: database of antimicrobial/cytotoxic activity and structure of peptides as a resource for development of new therapeutics. Nucleic Acids Res. 49, D288–D297. 10.1093/nar/gkaa991.

22. Wan, F., Torres, M.D.T., Peng, J., and de la Fuente-Nunez, C. (2024). Deep-learning-enabled antibiotic discovery through molecular de-extinction. Nat. Biomed. Eng. 8, 854–871. 10.1038/s41551-024-01201-x.

23. Özçelik, R., Brinkmann, H., Criscuolo, E., and Grisoni, F. (2025). Generative Deep Learning for de Novo Drug Design&#xe5f8;A Chemical Space Odyssey. J. Chem. Inf. Model. 65, 7352–7372. 10.1021/acs.jcim.5c00641.

24. Micsonai, A., Wien, F., Murvai, N., Nyiri, M.P., Balatoni, B., Lee, Y.-H., Molnár, T., Goto, Y., Jamme, F., and Kardos, J. (2025). BeStSel: analysis site for protein CD spectra—2025 update. Nucleic Acids Res. 53, W73–W83. 10.1093/nar/gkaf378.

25. Malanovic, N., and Lohner, K. (2016). Gram-positive bacterial cell envelopes: The impact on the activity of antimicrobial peptides. Biochim. Biophys. Acta BBA – Biomembr. 1858, 936–946. 10.1016/j.bbamem.2015.11.004.

26. Porto, W.F., Fensterseifer, I.C.M., Ribeiro, S.M., and Franco, O.L. (2018). Joker: An algorithm to insert patterns into sequences for designing antimicrobial peptides. Biochim. Biophys. Acta BBA – Gen. Subj. 1862, 2043–2052. 10.1016/j.bbagen.2018.06.011.

27. Zhang, M., Ouyang, J., Fu, L., Xu, C., Ge, Y., Sun, S., Li, X., Lai, S., Ke, H., Yuan, B., et al. (2022). Hydrophobicity Determines the Bacterial Killing Rate of α-Helical Antimicrobial Peptides and Influences the Bacterial Resistance Development. J. Med. Chem. 65, 14701–14720. 10.1021/acs.jmedchem.2c01238.

28. Richter, A., Sutherland, D., Ebrahimikondori, H., Babcock, A., Louie, N., Li, C., Coombe, L., Lin, D., Warren, R.L., Yanai, A., et al. (2022). Associating Biological Activity and Predicted Structure of Antimicrobial Peptides from Amphibians and Insects. Antibiotics 11, 1710. 10.3390/antibiotics11121710.

29. Eliseev, I.E., Terterov, I.N., Yudenko, A.N., and Shamova, O.V. (2019). Linking sequence patterns and functionality of alpha-helical antimicrobial peptides. Bioinformatics 35, 2713–2717. 10.1093/bioinformatics/bty1048.

30. Li, C., Sutherland, D., Richter, A., Coombe, L., Yanai, A., Warren, R.L., Kotkoff, M., Hof, F., Hoang, L.M.N., Helbing, C.C., et al. (2024). De novo synthetic antimicrobial peptide design with a recurrent neural network. Protein Sci. 33, e5088. 10.1002/pro.5088.

31. Baek, M.-H., Kamiya, M., Kushibiki, T., Nakazumi, T., Tomisawa, S., Abe, C., Kumaki, Y., Kikukawa, T., Demura, M., Kawano, K., et al. (2016). Lipopolysaccharide-bound structure of the antimicrobial peptide cecropin P1 determined by nuclear magnetic resonance spectroscopy. J. Pept. Sci. 22, 214–221. 10.1002/psc.2865.

32. Mohanram, H., and Bhattacharjya, S. (2016). ’Lollipop’-shaped helical structure of a hybrid antimicrobial peptide of temporin B-lipopolysaccharide binding motif and mapping cationic residues in antibacterial activity. Biochim. Biophys. Acta 1860, 1362–1372. 10.1016/j.bbagen.2016.03.025.

33. Bhunia, A., Mohanram, H., Domadia, P.N., Torres, J., and Bhattacharjya, S. (2009). Designed β-Boomerang Antiendotoxic and Antimicrobial Peptides: STRUCTURES AND ACTIVITIES IN LIPOPOLYSACCHARIDE*♦. J. Biol. Chem. 284, 21991–22004. 10.1074/jbc.M109.013573.

34. Bhunia, A., Domadia, P.N., Torres, J., Hallock, K.J., Ramamoorthy, A., and Bhattacharjya, S. (2010). NMR Structure of Pardaxin, a Pore-forming Antimicrobial Peptide, in Lipopolysaccharide Micelles: MECHANISM OF OUTER MEMBRANE PERMEABILIZATION 2. J. Biol. Chem. 285, 3883–3895. 10.1074/jbc.M109.065672.

35. Yagi-Utsumi, M., Yamaguchi, Y., Boonsri, P., Iguchi, T., Okemoto, K., Natori, S., and Kato, K. (2013). Stable isotope-assisted NMR characterization of interaction between lipid A and sarcotoxin IA, a cecropin-type antibacterial peptide. Biochem. Biophys. Res. Commun. 431, 136–140. 10.1016/j.bbrc.2013.01.009.

36. Hakoshima, T. (2005). Leucine Zippers. In Encyclopedia of Life Sciences (Wiley). 10.1038/npg.els.0005049.

37. Nang, S.C., Azad, M.A.K., Velkov, T., Zhou, Q. (Tony), and Li, J. (2021). Rescuing the Last-Line Polymyxins: Achievements and Challenges. Pharmacol. Rev. 73, 679–728. 10.1124/pharmrev.120.000020.

38. Ellenberger, T.E., Brandl, C.J., Struhl, K., and Harrison, S.C. (1992). The GCN4 basic region leucine zipper binds DNA as a dimer of uninterrupted α Helices: Crystal structure of the protein-DNA complex. Cell 71, 1223–1237. 10.1016/S0092-8674(05)80070-4.

39. Le, C.-F., Fang, C.-M., and Sekaran, S.D. (2017). Intracellular Targeting Mechanisms by Antimicrobial Peptides. Antimicrob. Agents Chemother. 61, e02340–16. 10.1128/AAC.02340-16.

40. Hancock, R.E.W., and Sahl, H.-G. (2006). Antimicrobial and host-defense peptides as new anti-infective therapeutic strategies. Nat. Biotechnol. 24, 1551–1557. 10.1038/nbt1267.

41. Moffatt, J.H., Harper, M., Harrison, P., Hale, J.D.F., Vinogradov, E., Seemann, T., Henry, R., Crane, B., St. Michael, F., Cox, A.D., et al. (2010). Colistin Resistance in Acinetobacter baumannii Is Mediated by Complete Loss of Lipopolysaccharide Production. Antimicrob. Agents Chemother. 54, 4971–4977. 10.1128/AAC.00834-10.

42. Campos, M.A., Vargas, M.A., Regueiro, V., Llompart, C.M., Albertí, S., and Bengoechea, J.A. (2004). Capsule Polysaccharide Mediates Bacterial Resistance to Antimicrobial Peptides. Infect. Immun. 72, 7107–7114. 10.1128/IAI.72.12.7107-7114.2004.

43. Peschel, A., and Sahl, H.-G. (2006). The co-evolution of host cationic antimicrobial peptides and microbial resistance. Nat. Rev. Microbiol. 4, 529–536. 10.1038/nrmicro1441.

44. Soares, D., Hetzel, L., Szymczak, P., Torres, M.D.T., Sommer, J., Fuente-Nunez, C. de la, Theis, F.J., Günnemann, S., and Szczurek, E. (2026). OmegAMP: Targeted AMP Discovery via Biologically Informed Generation. Trans. Mach. Learn. Res.

45. Veltri, D., Kamath, U., and Shehu, A. (2018). Deep learning improves antimicrobial peptide recognition. Bioinformatics 34, 2740–2747. 10.1093/bioinformatics/bty179.

46. Jhong, J.-H., Yao, L., Pang, Y., Li, Z., Chung, C.-R., Wang, R., Li, S., Li, W., Luo, M., Ma, R., et al. (2021). dbAMP 2.0: updated resource for antimicrobial peptides with an enhanced scanning method for genomic and proteomic data. Nucleic Acids Res. 50, D460–D470. 10.1093/nar/gkab1080.

47. Shi, G., Kang, X., Dong, F., Liu, Y., Zhu, N., Hu, Y., Xu, H., Lao, X., and Zheng, H. (2022). DRAMP 3.0: an enhanced comprehensive data repository of antimicrobial peptides. Nucleic Acids Res. 50, D488–D496. 10.1093/nar/gkab651.

48. Cabas-Mora, G., Daza, A., Soto-García, N., Garrido, V., Alvarez, D., Navarrete, M., Sarmiento-Varón, L., Sepúlveda Yañez, J.H., Davari, M.D., Cadet, F., et al. (2024). Peptipedia v2.0: a peptide sequence database and user-friendly web platform. A major update. Database J. Biol. Databases Curation 2024, baae113. 10.1093/database/baae113.

49. UniProt Consortium (2023). UniProt: the Universal Protein Knowledgebase in 2023. Nucleic Acids Res. 51, D523–D531. 10.1093/nar/gkac1052.

50. Wimley, W.C., and White, S.H. (1996). Experimentally determined hydrophobicity scale for proteins at membrane interfaces. Nat. Struct. Mol. Biol. 3, 842–848. 10.1038/nsb1096-842.

51. Dawson, R.M.C. (1986). Data for biochemical research 3rd ed. (Clarendon Press).

52. Levitt, M. (1978). Conformational preferences of amino acids in globular proteins. Biochemistry 17, 4277–4285. 10.1021/bi00613a026.

53. Zhao, G., and London, E. (2006). An amino acid “transmembrane tendency” scale that approaches the theoretical limit to accuracy for prediction of transmembrane helices: Relationship to biological hydrophobicity. Protein Sci. 15, 1987–2001. 10.1110/ps.062286306.

54. Juretić, D., Vukicević, D., Ilić, N., Antcheva, N., and Tossi, A. (2009). Computational design of highly selective antimicrobial peptides. J. Chem. Inf. Model. 49, 2873–2882. 10.1021/ci900327a.

55. Ronneberger, O., Fischer, P., and Brox, T. (2015). U-Net: Convolutional Networks for Biomedical Image Segmentation. Preprint at arXiv, 10.48550/arXiv.1505.04597 https://doi.org/10.48550/arXiv.1505.04597.

56. Ho, J., Jain, A., and Abbeel, P. (2020). Denoising Diffusion Probabilistic Models. Preprint at arXiv, 10.48550/arXiv.2006.11239 https://doi.org/10.48550/arXiv.2006.11239.

57. Nichol, A., and Dhariwal, P. (2021). Improved Denoising Diffusion Probabilistic Models. Preprint at arXiv, 10.48550/arXiv.2102.09672 https://doi.org/10.48550/arXiv.2102.09672.

58. Salimans, T., and Ho, J. (2022). Progressive Distillation for Fast Sampling of Diffusion Models. Preprint at arXiv, 10.48550/arXiv.2202.00512 https://doi.org/10.48550/arXiv.2202.00512.

59. Conchillo-Solé, O., de Groot, N.S., Avilés, F.X., Vendrell, J., Daura, X., and Ventura, S. (2007). AGGRESCAN: a server for the prediction and evaluation of “hot spots” of aggregation in polypeptides. BMC Bioinformatics 8, 65. 10.1186/1471-2105-8-65.

60. Wang, R., Wang, T., Zhuo, L., Wei, J., Fu, X., Zou, Q., and Yao, X. (2024). Diff-AMP: tailored designed antimicrobial peptide framework with all-in-one generation, identification, prediction and optimization. Brief. Bioinform. 25, bbae078. 10.1093/bib/bbae078.

61. Müller, A.T., Hiss, J.A., and Schneider, G. (2018). Recurrent Neural Network Model for Constructive Peptide Design. J. Chem. Inf. Model. 58, 472–479. 10.1021/acs.jcim.7b00414.

62. Nagarajan, D., Nagarajan, T., Roy, N., Kulkarni, O., Ravichandran, S., Mishra, M., Chakravortty, D., and Chandra, N. (2018). Computational antimicrobial peptide design and evaluation against multidrug-resistant clinical isolates of bacteria. J. Biol. Chem. 293, 3492–3509. 10.1074/jbc.M117.805499.

63. Luo, Z., Geng, A., Wei, L., Zou, Q., Cui, F., and Zhang, Z. (2025). CPL-Diff: A Diffusion Model for De Novo Design of Functional Peptide Sequences with Fixed Length. Adv. Sci. 12, 2412926. 10.1002/advs.202412926.

64. Steinegger, M., and Söding, J. (2017). MMseqs2 enables sensitive protein sequence searching for the analysis of massive data sets. Nat. Biotechnol. 35, 1026–1028. 10.1038/nbt.3988.

65. Nell, M.J., Tjabringa, G.S., Wafelman, A.R., Verrijk, R., Hiemstra, P.S., Drijfhout, J.W., and Grote, J.J. (2006). Development of novel LL-37 derived antimicrobial peptides with LPS and LTA neutralizing and antimicrobial activities for therapeutic application. Peptides 27, 649–660. 10.1016/j.peptides.2005.09.016.

